# Integrating Machine Learning with Flow-Imaging Microscopy for Automated Monitoring of Algal Blooms

**DOI:** 10.1101/2024.11.12.623192

**Authors:** Farhan Khan, Benjamin Gincley, Andrea Busch, Dienye L Tolofari, John W Norton, Emily Varga, R Michael Mckay, Miguel Fuentes-Cabrera, Tad Slawecki, Ameet J. Pinto

## Abstract

Real-time monitoring of phytoplankton in freshwater systems is critical for early detection of harmful algal blooms so as to enable efficient response by water management agencies. This paper presents an image processing pipeline developed to adapt ARTiMiS, a low-cost automated flow-imaging device, for real-time algal monitoring specifically in freshwater and environmental systems. This pipeline addresses several challenges associated with autonomous imaging of aquatic samples such as flow-imaging artifacts (i.e., out-of-focus and background objects), as well as specific challenges associated with monitoring of open environmental systems (i.e., identification of novel objects). The pipeline leverages a Random Forest model to identify out-of-focus particles with an accuracy of 89% and a custom background particle detection algorithm to identify and remove particles that erroneously appear in consecutive images with >97±2.8% accuracy. Furthermore, a convolutional neural network (CNN), trained to classify distinct classes comprising both taxonomical and morphological categories, achieved 94% accuracy in a closed dataset. Nonetheless, the supervised closed-set classifiers struggled with the accurate classification of objects when challenged with debris and novel particles which are common in complex open environments; this limits real-time monitoring applications by requiring extensive manual oversight. To mitigate this, three methods incorporating classification with rejection were tested to improve model precision by excluding irrelevant or unknown classes. Combined, these advances present a fully integrated, end-to-end solution for real-time HAB monitoring in open environmental systems thus enhancing the scalability of automated detection in dynamic aquatic environments.

**Highlights:**

- Random Forest model is more generalizable than Convolutional Neural Networks to remove out-of-focus particles.
- A two-stage clustering algorithm is effective at removing background particles in flow imaging microscopy.
- Closed-set CNN classifier performance deteriorates when challenged with unknown particles.
- Classification with rejection improves both precision and accuracy for environmental samples.

## Introduction

Increasing temperatures, widespread fertilizer application, and consequent nutrient loading into water bodies result in excessive phytoplankton growth, including cyanobacterial harmful algal blooms (cyanoHABs). CyanoHABs are associated with toxin release and can also cause water discoloration and taste and odor problems in the drinking water supply (Cen et al., 2020; Jetoo et al., 2015; Kramer et al., 2018; Watson et al., 2008; Zhang et al., 2011). Toxicity and hypoxia caused by these blooms can result in mortality to wildlife, loss of habitat for aquatic life, and economic losses for tourism and fisheries industries (Dodds et al., 2009; Heil and Muni-Morgan, 2021; Smith et al., 2019). CyanoHAB monitoring is currently performed using manual microscopy, fluorometry, molecular assays, or remote sensing (Caballero et al., 2020; Den Uyl et al., 2022; Richardson et al., 2010; Sanseverino et al., 2017; Stauffer et al., 2019; Stroming et al., 2020; Toldrà et al., 2019). While manual microscopy offers detailed taxonomic information, it is time-consuming and requires expert taxonomists. Fluorescence probes and remote sensing can estimate phytoplankton biomass in bulk but lack the taxonomic detail necessary for actionable insights. Specifically, this taxonomic information is crucial for managing blooms and determining the appropriate response, as specific phytoplankton species are responsible for toxin production, taste and odor issues, and filter clogging at drinking water treatment plants (DWTPs). Therefore, there is a pressing need for an automated imaging approach that is high-throughput and capable of providing quantitative taxonomic information.

There have been important advances in both imaging hardware and software such that distributed and automated real-time imaging for quantitative taxonomic data is likely feasible. Specifically, low-cost field deployable imaging solutions like LudusScope (Kim et al., 2016), SAMSON (Deglint et al., 2018), HABscope (Hardison et al., 2019), PlanktoScope (Pollina et al., 2022), and ARTiMiS (Gincley et al., 2024) are available as alternatives to more expensive benchtop phytoplankton imaging platforms like FlowCam (Sieracki et al., 1998) and CytoSense XR (CytoBuoy) (Fragoso et al., 2019). Similarly, there have been significant advances in image processing as well. Existing commercial software, such as Visual Spreadsheet, uses statistical filters based on morphological properties to sort phytoplankton into different classification bins. However, this approach struggles to differentiate between phytoplankton with similar features (e.g., size) (Camoying and Yñiguez, 2016; Owen et al., 2022). In contrast, advances in computing power, the availability of large datasets, and the popularization of Convolutional Neural Networks (CNNs) have revolutionized image-processing approaches. This shift has popularized phytoplankton classification and significantly increased its accuracy (Eerola et al., 2024). Research on phytoplankton classification surged over the last decade, especially since 2014 (Ciranni et al., 2024). Recently, Eerola et al. (2024) provided a comprehensive overview of research on automated plankton classification and identified some key associated challenges.

Despite these advancements and the high classification accuracies reported in the literature, widespread automated plankton monitoring has not yet been realized. This is largely because a holistic solution—integrating both hardware and software—that can process environmental samples and provide taxonomy-based quantitative results on phytoplankton communities has yet to be developed. In addition to classification challenges, automated phytoplankton monitoring using flow imaging microscopy struggles with accurate classification of images acquired under field-relevant conditions which often include out-of-focus particles, background particles, abiotic particles, and out-of-distribution particles. Out-of-focus particles (OOF) are a very common issue with flow imaging microscopy (Camoying and Yñiguez, 2016). Similarly, it is not uncommon for particles to get stuck in imaging flow channels and appear in multiple frames (i.e., background particles) resulting in duplicate counting (Álvarez et al., 2012; Zölls et al., 2013). Often, manual processing is required to exclude OOF or background particles (Álvarez et al., 2012; Romero-Martínez et al., 2017) from images; this is not consistent with the goal of automated imaging and image analyses. Out-of-distribution (OOD) particles are particles that lack a representative image or image collection (non-annotated and previously unseen plankton and non-plankton particles) in the training phase of model construction. Open aquatic systems are hyper-diverse. It is unrealistic to expect, at least currently, that the imaging platform will never encounter an OOD particle. Traditional CNN-based classifiers tend to misclassify these novel particles, making the classification process unreliable and again, necessitating manual curation. The requirement for such post-processing has hindered the widespread adoption of flow imaging microscopy for automated environmental monitoring.

This study focuses on the development of an image-processing pipeline that addresses some of the key challenges hindering automated plankton monitoring. Here, a low-cost miniaturized flow imaging microscope (FIM), i.e., ARTiMiS (Gincley et al., 2024), was used for sample processing and image collection. ARTiMiS was developed to make microalgal monitoring more affordable and accessible to a broader range of end users. Gincley et al. (2024) demonstrated that this device and its image-processing workflow can be used for real-time microalgal monitoring in engineered systems designed for specific microalgae (i.e., low likelihood of encountering novel particles). In this study, the ARTiMiS was adapted for phytoplankton monitoring in environmental systems, and an image processing pipeline was designed and developed to address challenges associated with environmental samples, reducing the need for manual intervention. Although this pipeline was specifically developed for ARTiMiS, the steps can be implemented for other automated image acquisition platforms.

## Materials and Methods

### ARTiMiS Flow Imaging Microscope

ARTiMiS (Gincley et al., 2024) is a low-cost self-contained autonomous microscope that provides 5X magnification onto a Raspberry Pi Camera V2 (Sony IMX219 sensor) with a sample imaging resolution of 1.55 µm, enabling detailed imaging suitable for environmental monitoring applications. At this level of magnification, a 5 µm particle is represented by a 29.5-pixel image, providing sufficient detail for accurate classification and analysis of microscopic particles. Illumination is provided by a programmable LED array enabling both bright and dark field illumination. An aqueous sample is circulated through a microfluidic chip with a 200 µm depth flow channel via a stop-flow mechanism. The device is equipped with autofocusing as an automatic routine before and during sample processing. A graphical user interface allows users to set parameters for a sample run including settling time, illumination mode, number of images to be captured, autofocusing frequency, etc. See (Gincley et al., 2024) for a complete description of the instrument.

### Collection of Plankton Images

Samples were collected by the Great Lakes Water Authority (GLWA) from the intake point of Lake Huron Water Treatment Facility, Water Works Park, and Southwest Water Treatment Plant and shipped overnight to Georgia Institute of Technology weekly during summer months (May-Sep) and bi-weekly basis during winter months (Oct-Apr) from April 2022 to November 2023. Particle concentrations varied seasonally and were below the detection limit of the ARTiMiS (∼3.2 x 10^3^ particles/mL) (Gincley et al., 2024) during April and May. To address this issue, samples were concentrated using a plankton net for 30 minutes at a flow rate of 6.8 liters/minute. Then the concentrate was eluted in 50 mL of water and prepared for shipping. Samples were also collected from Lake St. Clair at the Stoney Point Water Treatment Plant, Stoney Point, Ontario, Canada. These samples were collected from a tap installed at the intake well and concentrated from 218 L to 50 mL using a plankton net. In addition to the field samples, *Aphanizomenon sp*. (CAWBG01), *Planktothrix agardhii*, and *Dolichospermum flos-aquae* were cultured in the laboratory to generate a collection of high-priority cyanobacterial species impacting the Great Lakes region. These cultures were maintained in Jaworski medium at 25°C under a 12h:12h day-night cycle at a photosynthetic active radiation (PAR) value of 80 µmol-photons m^-2^·s^-1^. Cultures were grown in polycarbonate, unbaffled, sterile vent-cap flasks at nominal flask volumes of 125 mL (VWR International, 89095-258) and were agitated continuously using an orbital shaker. Field samples and laboratory cultures were imaged using ARTiMiS under both bright-field and dark-field modalities. A total of 30 to 90 images were collected for each sample, with a settling time ranging from 60 to 90 seconds before imaging. Auto-focus was enabled for high-density samples and disabled for low-density samples.

### Development of the Image Processing Workflow

Figure 1 provides a complete overview of the developed image processing workflow and subsequent sections provide details for each step.

**Figure 1:**
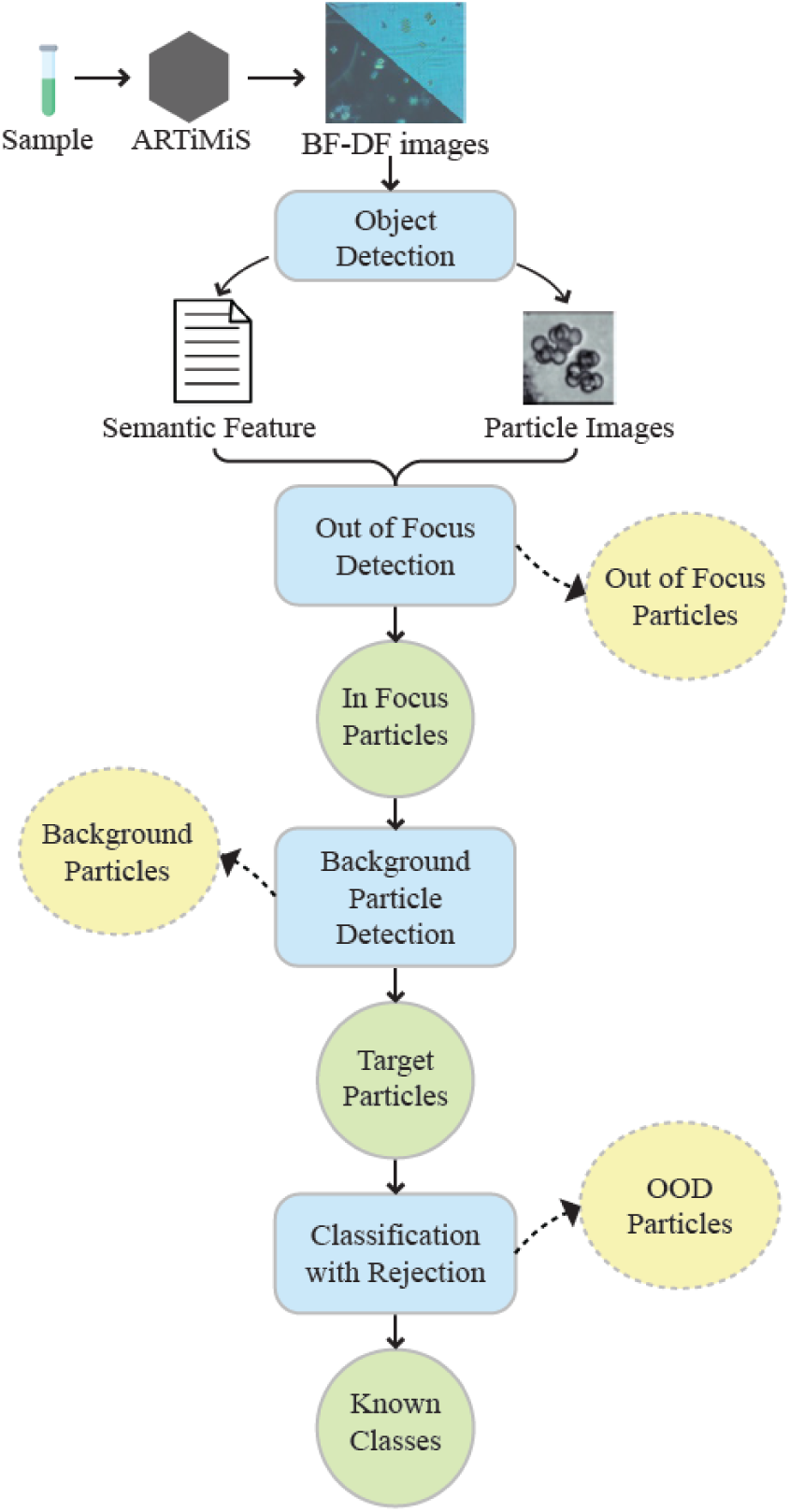
Schematic of image processing pipeline. Brightfield and darkfield images are collected using ARTiMiS. Object detection algorithm finds particles and calculates semantic features of each particle. These particle images and semantic features are processed for out-of-focus particle designation, background particle detection, and classification with rejection steps sequentially, thus discarding out-of-focus particles, background particles and out-of-distribution (OOD) particles in the process. As a result of this processing, labels are assigned for each particle observed in the raw image dataset to be used in aggregate analysis.

### Object Detection

Widefield images collected during the sample run were processed using an object detection algorithm (ODA) to detect particles and extract their cropped images. Here, a “particle” refers to any contiguous object detected within a specified region of interest (ROI), allowing for precise segmentation and examination of individual entities. Gincley et al. (2024) developed two ODAs for the ARTiMiS: of these, the Segmenter ODA was better suited for the variable particle sizes typically encountered in environmental samples and thus, it was selected for all image processing. Pixel coordinates and 47 geometric features were calculated for each detected particle. A description of the geometric features can be found in the Supplementary Information (**Table S1**).

### OOF Particle Identification

An OOF particle classifier was integrated into the image processing pipeline to identify and exclude OOF particles from further analysis. This was a Random Forest (RF) classifier trained using the geometric features of the particles. For training and testing, a total of 3,470 in-focus (IF) particles and 2,663 OOF particles were manually annotated and then divided into train and test sets at a 70:30 ratio. Pearson correlation analysis was conducted to detect and remove features with high autocorrelation, resulting in the elimination of 14 features. This resulted in the use of 33 features, which were used to train an RF model using the scikit-learn library in Python (v1.1.1). The features were then ranked in order of importance for class identity (Gincley et al., 2024). Following this analysis, the model was retrained using the 12 top-ranked features to optimize its accuracy.

### Background Particle Detection

A two-stage background particle detection algorithm was developed to detect particles stuck in the field of view (FOV) in consecutive image frames. This algorithm clusters particles in three consecutive image frames based on their x-y pixel coordinates. Pairwise distances are calculated for every particle in the three consecutive frames. Particles with pairwise distances smaller than a user-defined threshold are clustered using the Density-Based Spatial Clustering of Applications with Noise (DBSCAN) method (Ester et al., 1996). A moving particle can sometimes be imaged within the defined threshold of a stationary particle. To address this, particles within a cluster based on the pairwise distance were grouped into subclusters based on their features. This resulted in subclusters consisting of particles that appeared at similar locations in the wide-field images and with similar, if not identical, features. Particles within subclusters were flagged as background particles, while particles outside of clusters were considered non-background or valid particles. In theory, this two-stage clustering method should be able to detect whether the same particle appears in two consecutive images. However, smaller debris may occasionally appear in the same ROIs, leading to measurement variation for the same particle. This can lead to incorrect identification of background particles: the particle with debris along with its immediate previous and next particles might be misclassified. To reduce misclassification, the algorithm flagged a particle as a background particle only when it was identified twice in three consecutive frames.

### Convolution Neural Networks

CNN architectures employed for the image classification task were based on the “micro-CNN” framework introduced previously (Gincley et al., 2024). For this work, the CNN architecture was optimized through iterative evaluation of convolution layer configurations and dropout rates. The feature extraction pipeline was constructed using a sequence of convolutional layers, each followed by max-pooling and dropout operations.

Configurations involving three, four, and five layers were explored to ascertain the optimal depth of the network. Convolution kernel sizes were also modified and evaluated. To ensure that a model was not overfitted to the training dataset, varying dropout values were tested (Srivastava et al., 2014). After extensive hyperparameter tuning, the final model architecture was established as one block of CONV-MP, four sequential blocks of CONV-MP-DO, followed by a fully connected (dense) layer prior to model output. In this configuration, “CONV” describes a 2D convolutional layer, “MP” denotes a max pooling layer, and “DO” represents a dropout layer. The model employed a SoftMax activation function for output classification, and a dropout rate of 0.30 was used. During training, the Adam optimizer was used with a 0.001 learning rate and categorical cross-entropy as the loss function. The model configuration that provided the highest overall accuracy while reducing inter-class accuracy disparity was chosen for this study. A similar architecture was used for OOD identification. The structure of the OOD classifier was a variation of the described model, also using SoftMax as the output activation function and trained with the Adam optimizer (0.001 learning rate, categorical cross-entropy).

### OOD Particle Identification

The SoftMax function in the final layer of a CNN generates probability scores for each class. The SoftMax thresholding method operates under the assumption that SoftMax scores for in-distribution (known) class images are high, whereas scores for ambiguous or OOD images are low. By establishing a cutoff score (threshold), it is possible to differentiate between known and OOD particles. In this study, particle images with SoftMax scores falling below the threshold were classified as OOD particles. The threshold was optimized based on the highest accuracy achieved during evaluation (see results and discussion). Monte Carlo dropout (MCD), while sharing similarities with SoftMax thresholding in calculating probability scores, employs a distinct approach. Unlike SoftMax thresholding, MCD involves performing multiple forward passes through the network with varied dropout configurations to estimate model uncertainty. This variability in probability scores across different network configurations can assist in distinguishing OOD particles. For each particle image, 50 forward passes were executed, and the mean probability score was computed. The threshold for identifying OOD particles was then determined using the same methodology as with SoftMax thresholding. The third method, Class Anchor Clustering (Miller et al., 2021), leverages a distance-based loss function. This loss function trains the classification model to minimize intra- class distance while maximizing inter-class distance. Theoretically, this should enhance the separation of OOD particles from the known classes by creating more distinct class boundaries.

### Train-Test Dataset

Given the high observed particle diversity, a combination of taxonomy and morphotype labels was employed for image annotation. Images were categorized into 12 distinct classes, with taxonomic labels including *Microcystis*, *Merismopedia*, *Staurastrum*, and *Fragilaria*. Morphotype labels included descriptors like small cells, single round cells, and filaments. Employing this hybrid approach, a total of 11,000 particle images were manually annotated from field samples and laboratory cultures. From the 11,000 annotated particle images, 5,000 particle images across 12 classes were used for training, validation, and testing of image classification models. Examples of these classes are shown in **Figure S1**. Each of these classes contained at least 60 images, with the number of unique objects per class detailed in **Table S2**.

Images from each class were divided into three subsets: train set, validation set, and test set in a 70:15:15 ratio. To enhance model generalization and increase the diversity of the training data, each image in the training set underwent augmentation. This augmentation was implemented as a set of 90° rotations and mirrors, resulting in 8 non-destructive transformations for each original image. This approach effectively expanded the training set and assisted in the learning of rotational invariance and feature robustness, improving model performance in classification tasks. The training and validation sets were used for model development, while the unseen test set was reserved for evaluating performance once training was complete.

## Results and Discussion

### Ecosystem-specific calibration improves the performance of ODA for freshwater phytoplankton imaging

Gincley et al. (2024) demonstrated that the “Segmenter” object detection algorithm (ODA) outperformed the “Fast Object Detector” in correctly processing images containing particles of varying sizes and morphologies. In this study, the efficacy of the Segmenter algorithm was evaluated for accurately detecting phytoplankton in images from freshwater samples collected in the Great Lakes region. The Segmenter algorithm was initially calibrated for algal communities in a full-scale microalgal cultivation system designed for nutrient recovery from wastewater.

While the algorithm performed well for the majority of imaged particles, it encountered challenges with large colonial morphologies, resulting in the erroneous splitting of colonies into multiple objects. To address this limitation, the Segmenter algorithm was calibrated by adjusting its parameters to better accommodate the diverse cell types, sizes, and morphologies characteristic of freshwater samples. The object detection performance of the newly calibrated ODA was compared to the baseline performance (prior calibration) on samples with varying particle concentration (i.e., particles per unit volume) and FOV coverage; here, FOV coverage refers to the percent of the surface area in FOV that is occupied by particles. A dense (1.65x10^6^ particle/ mL or 2.8% FOV coverage) and a sparse (4x10^5^ particle/mL or 0.7% FOV coverage) sample were chosen as such they closely resemble the density of samples (“EcoRecover medium density” (Eco Med) and “EcoRecover high density” (Eco High)) from the referenced dataset used in the prior calibration. Multiple images for dense and sparse samples were selected and objects in these images were annotated manually. For evaluation, each ROI identified by the ODA was classified into five categories: true positive (i.e., single particle in ROI), false positive (i.e., no particle in ROI), false negative (i.e., particle not entirely in ROI), merge (i.e., more than one particle in one ROI), and split (i.e., one particle distributed over more than ROI).

The calibrated algorithm showed similar performance to the baseline algorithm for sparse samples. The particle size distribution in sparse samples closely resembles that of the Eco Med sample (Figure 2A **and 2B**). This suggested that the performance of the calibrated model was as good as the baseline for particle distributions similar to EcoRecover samples.

**Figure 2:**
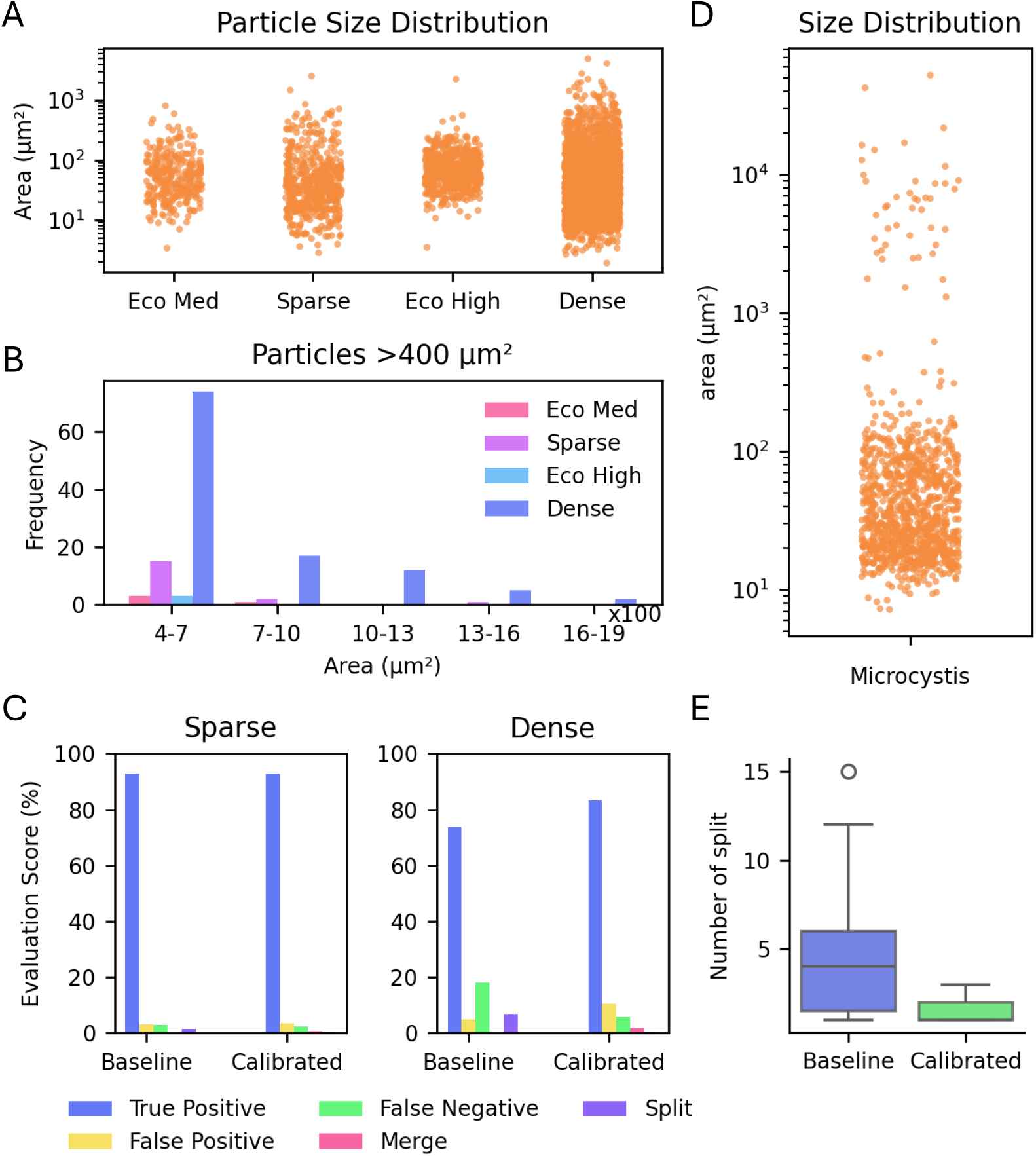
Particle size distribution and evaluation of particle identification for baseline vs. calibrated ODA. (A) Particle size distribution in EcoRecover medium and high-density samples and freshwater sparse and dense samples. (B) Histogram of particles with an area >400 µm² in EcoRecover medium and high density and freshwater sparse and dense sample. (C) Performance comparison of baseline and calibrated ODA for sparse and dense samples. (D) Particle size distribution for Microcystis colonies. (E) Number of splits performed by baseline and calibrated ODA for Microcystis colonies

Dense freshwater samples contained relatively higher colonial or large particles (area >400 µm²) compared to the original dataset (Figure 2B). Calibrated ODA achieved 83% accuracy for the dense sample whereas the baseline model exhibited 73.7% accuracy (Figure 2C). Calibrated ODA had more false positive incidents; however, among error types, false positives are of least concern given they can be identified as noise in downstream processing and discarded. On the other hand, calibrated ODA outperformed the reference baseline in terms of false negatives and split cases, an important reduction in error as these ROIs would otherwise be lost from analysis. Considering freshwater monitoring applications can encounter rare but impactful events, false negatives may lead to the failure to detect these occurrences. Surface water samples, including those in this study, often contain large plankton and colonies of plankton which are more susceptible to splitting during object detection. Thus, in the context of freshwater algae and cyanobacteria monitoring, minimizing the occurrences of false negatives and split cases is critical; the calibrated ODA performed similarly or better on these metrics (**Figures S2A-D**). The calibrated ODA was also tested on samples containing *Microcystis* colonies. Microcystis colony sizes can vary wildly, from 10 µm^2^ to more than 10^4^ µm^2^ (Figure 2D). When tested, the baseline ODA frequently splits these colonies into multiple smaller sections, or in some cases into individual cells, resulting in an over-enumeration of unique Microcystis colonies (**Figures S2E- F**). The calibrated ODA exhibited a much higher incidence of whole, intact colonial particle detection. Figure 2E highlights the split rate for each Microcystis colony by the two algorithms.

### Detection of OOF and background particles is critical for ensuring accurate quantitative monitoring

Visual artifacts such as OOF and background particles are inherent to flow-imaging microscopy (Choran and Örmeci, 2023; Marvin et al., 2019; Olsen and Adrian, 2000; Sediq et al., 2018; Zölls et al., 2013). Following object detection, the pool of candidate ROIs to be identified contains both in- and out-of-focus (IF, OOF) particles. Examples of IF and OOF can be found in **Figure S3**. To identify and exclude these OOF particles from downstream processing, a pre-processing step was developed. A Random Forest (RF) classification model was trained on 33 semantic features of the IF and OOF particles (collected during 2022 sampling period), achieving 90% overall classification accuracy. Feature importance analysis (Figure 3A) revealed that parameters associated with edge noise and luminance intensity were most effective at describing distinct differences between the two classes. Feature selection was performed iteratively to maximize the inclusion of informative features and minimize the inclusion of features irrelevant to accurate class discrimination. The incorporation of 12 top-ranked features resulted in the highest overall accuracy (91%) with 92% and 90% correct prediction for IF and OOF particles respectively (**Figure S5A**). Figure 3B shows the density plot for the top four features, while density plots for all 12 features are provided in **Figure S4**. Some separation between IF and OOF particle populations can be observed. As expected, misclassification predictions appear in the transition region of IF and OOF particles where the feature distributions of the two classes overlap. A suspected cause for this is due to the inherent nature of degree-of-focus as being a continuous quality; the binary nature of the classification scheme necessarily imposes a cutoff threshold within the overlapping region, and complete accuracy is not achievable.

**Figure 3:**
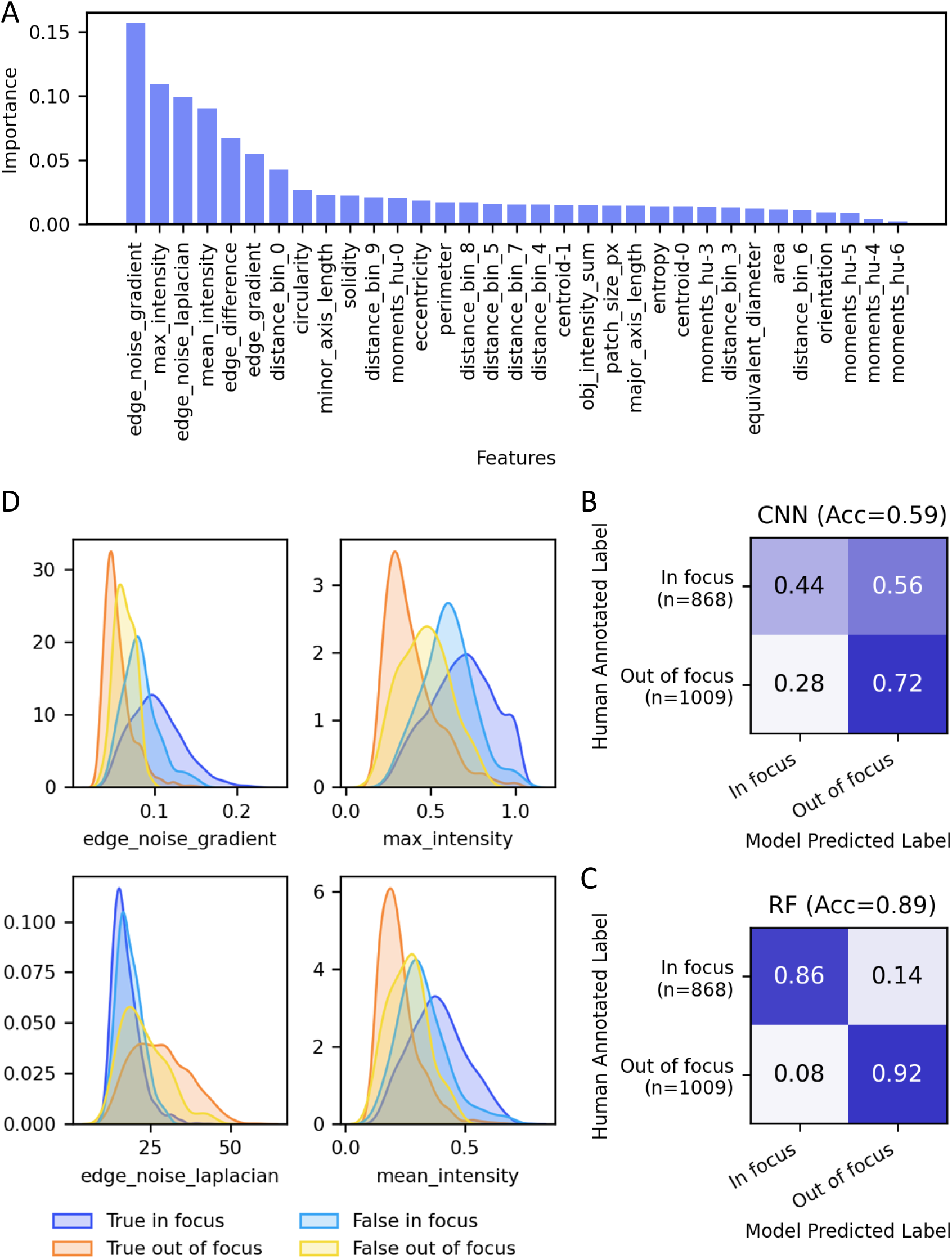
(A) Feature importance of the top 33 features relevant to in-focus (IF) and out-of-focus (OOF) classification from the random forest (RF) classifier. (B) and (C) Confusion matrices showing the classification performance of the RF and CNN models in identifying IF and OOF particles. (D) Density plot of the top four features i.e. edge noise gradient, max intensity, edge noise Laplacian, and mean intensity for true IF, true OOF, false IF, and false OOF particles.

In complement to the RF model, CNN was trained on the ROIs containing both IF and OOF particles with the aim of investigating whether a more complex model such as CNN could potentially enhance accuracy by leveraging image-based feature extraction. The CNN model achieved almost identical results (92% accuracy) to the RF model (**Figure S5B**). However, when the model was tested on sample data collected the following year, it exhibited a high misclassification rate (only 59% accuracy, Figure 3C). In contrast, the RF classifier achieved 89% accuracy (Figure 3D) on the same dataset, consistent with its performance on the dataset from the prior year (**Figure S5A).** This suggested that the RF model is more generalized compared to the CNN model. Therefore, the RF model was chosen for OOF identification for analysis of the complete sampling campaign.

In addition to OOF identification, background particle detection is important for real-time continuous monitoring of HABs, especially for low-density samples. Due to the low concentration of particles of interest, even a small number of persistent background particles can significantly skew results. A naïve approach involves capturing blank images before and after a sample run to identify background particles; however, this method is unreliable since blank images with ultrapure water can also contain contaminating particles, leading to erroneous background particle counts (Romero-Martínez et al., 2017). A number of dynamic scenarios may also occur: particles may get stuck in the field of view (FOV) mid-sample run, previously-stuck particles may dislodge at a later time, or a stagnant particle may experience a changed degree of focus due to mid-run autofocusing; all of these scenarios can result in background de-calibration. Thus, a static blank subtraction approach is not appropriate; instead, a dynamic approach is required to identify and remove background particles on a frame-by-frame basis. In theory, a background particle should appear in the same position in consecutive images. However, in a flow-through system, stagnant particles exhibit small shifts in position within the FOV during instrument operation as a result of fluid effects, small changes in camera or flow cell position, etc. These movements can also result in slight changes in the measured particle features. Successful removal of these stagnant background particles requires an approach that is tolerant to these perturbations in measurements.

A novel algorithm for background particle detection (BPD) was developed that tracks particles in successive frames based on their location and measured semantic features. The BPD algorithm was optimized for natural environmental samples and its performance was evaluated for low (1.6 x 10^5^ particles/mL), medium (3 x 10^5^ particles/mL), and medium-high (6 x 10^5^ particles/mL,) density samples. First, we identified 300 instances of stagnant particles and their corresponding coordinates in images collected from the select sample runs. Travel distances were calculated for these particles from one frame to the next frame using their x-y coordinates. Mean and median travel distances were found to be 16.8 and 13.15 pixels, respectively, with a standard deviation of 11.5 pixels (**Figure S6**). The tolerance of the BPD algorithm was calibrated to detect stagnant particles with >95% accuracy even when the particles may have moved by up to 20 pixels (Figure 4A). Subsequently, the calibrated algorithm was used on images from sample runs (real images) as well as synthetic images. Synthetic images were created by seeding blank images with both background particles and target particles. To simulate low-, medium- and high-density samples, blank images were seeded with combinations of 10, 15, and 25 target particles and 3, 5, and 8 background particles, respectively. These incidence rates corresponded to 0.3%, 0.5%, and 0.8% FOV coverage and sample concentrations of 2 x 10^5^, 3 x 10^5^, and 5 x 10^5^ particles per mL, respectively. Target particles to seed synthetic images were randomly selected from the annotated ROI image library and placed at random x-y coordinates in each frame. For background particle seeding, identical copies of a random particle image were placed at randomly chosen positions on the images. Positions of the background particles were randomly jittered no greater than 12 pixels from frame to frame. On synthetic images, BPD achieved >99% accuracy with a small number of false positives and achieved 97±2.8% accuracy for real images (from sample runs) with a small number of false negatives (Figure 4B **and S7**).

**Figure 4:**
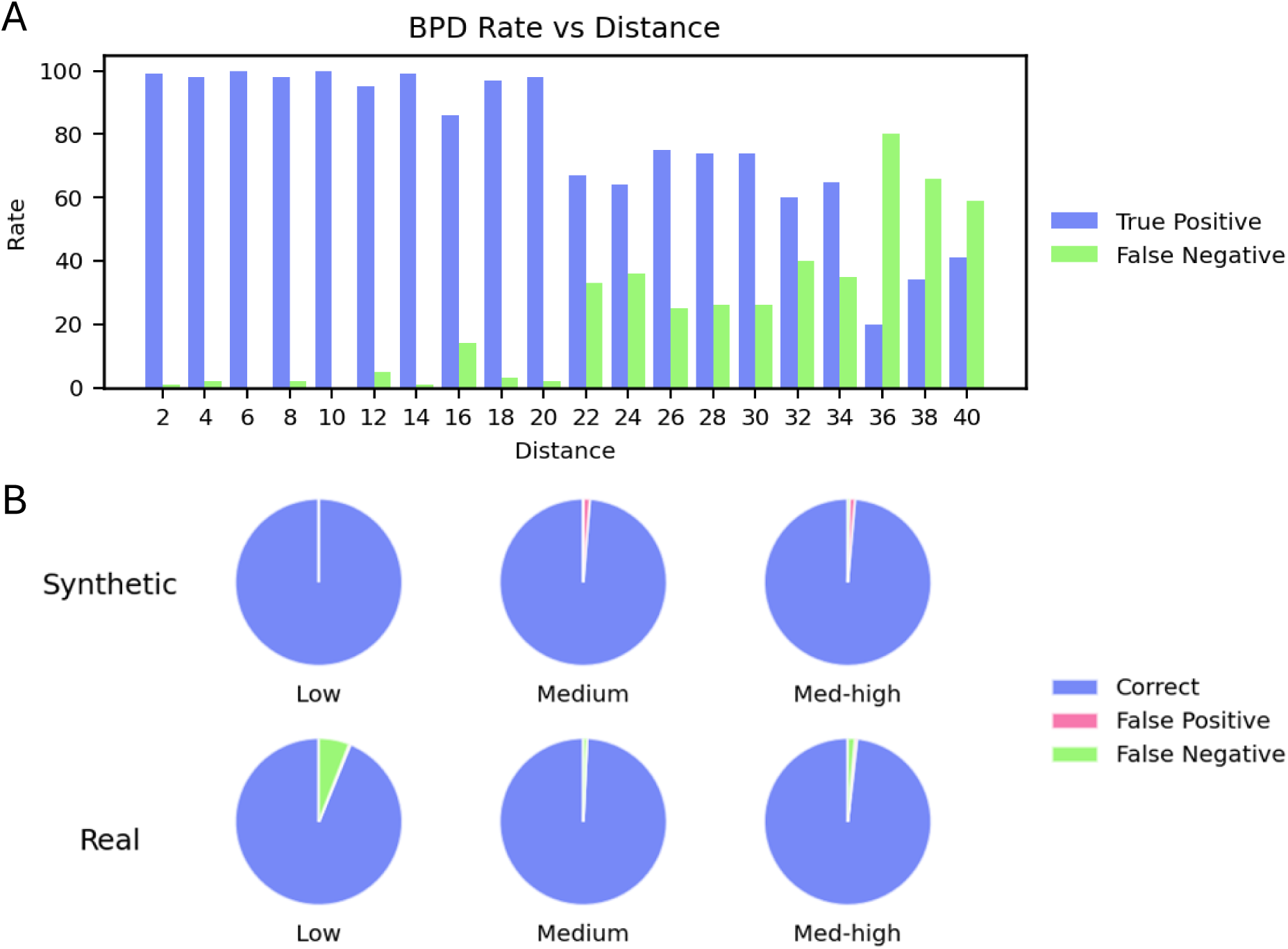
(A) Detection rate of background particles by the BPD algorithm at varying particle-to-particle distances. (B) Performance of the BPD algorithm on low-, medium-, and high-density real and synthetic samples.

### Closed-set classification accurately identifies classes included in the training dataset, but performance is degraded for OOD particle types

Previously, Gincley et. al. (2024) demonstrated that ARTiMiS could accurately classify particles from complex microalgal communities in a photobioreactor system and demonstrated that CNN models outperformed RF models for multiclass algal classification. Building on these findings, this study explored the application of CNN models for phytoplankton classification in freshwater samples. A custom CNN model was trained on 12 annotated classes, with the annotated dataset split for training, validation, and testing. The optimized CNN model achieved an overall accuracy of 94%, with ten classes exceeding 90% accuracy (Figure 5). The lower accuracy rates (mean 91.5% ±4.4%) were associated with colonial particles as compared to the single cell classes (mean 94.6% ± 5.7%). It is suspected that this disparity is due to the significant variation in size and shape of phytoplankton colonies, making their morphology challenging to generalize for a classification model.

**Figure 5:**
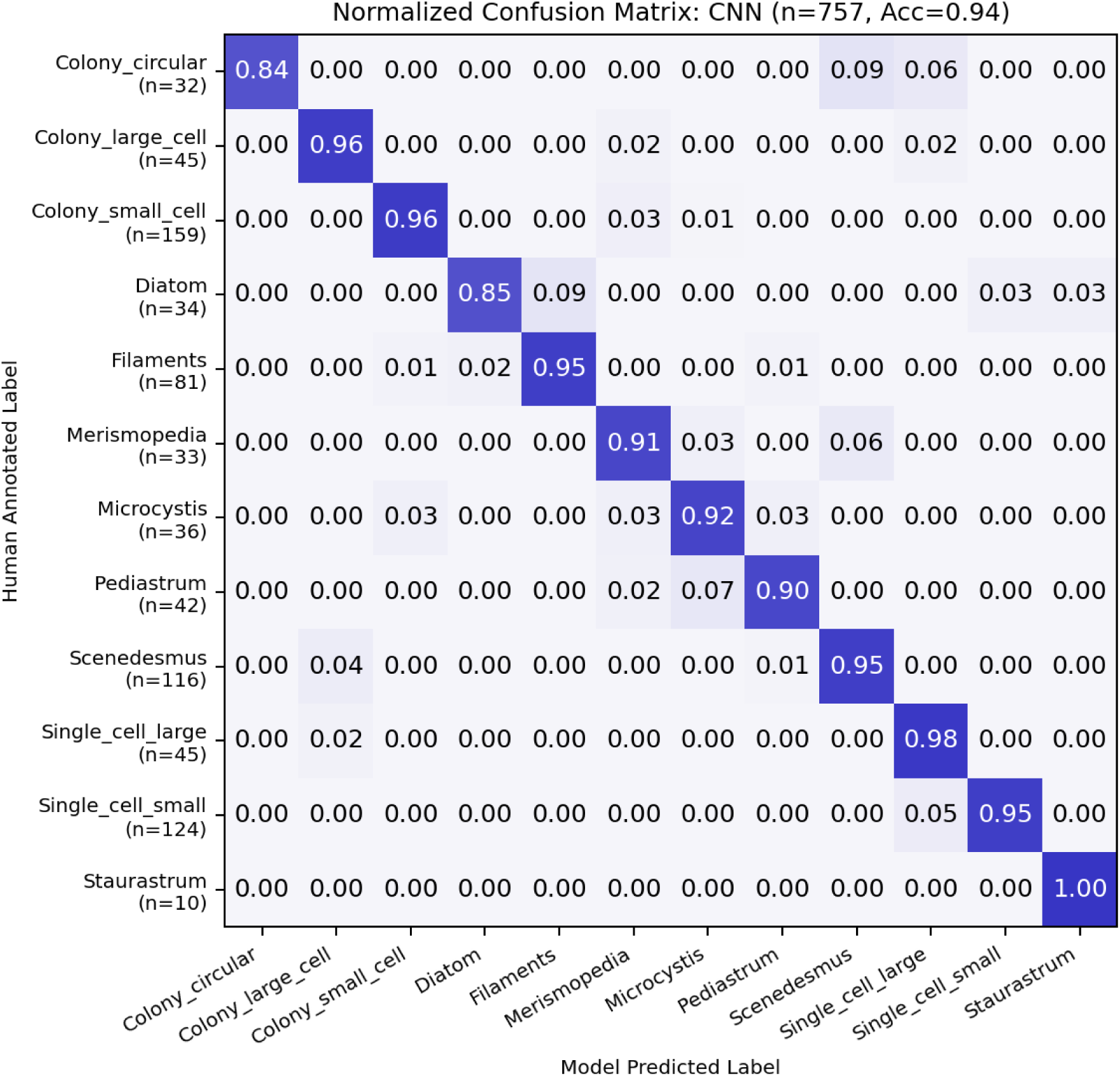
Confusion matrix showing performance of CNN model trained to distinguish among 12 classes of objects captured from environmental samples. Model accuracy, 94%, when evaluated on unseen annotated test data (n=757 particle images)

A considerable proportion of phytoplankton classification studies benchmark machine- and deep- learning model performance on standardized public datasets, e.g., WHOI dataset, ZOOSCAN dataset, ISIIS PlanktonSet-1.0 (Faillettaz et al., 2016; Heidi M. Sosik et al., 2015; Lumini and Nanni, 2019; Luo et al., 2018), or use meticulously curated internally maintained datasets (Chan et al., 2023; Correa et al., 2017; Xu et al., 2022). The most common class configuration is that of a “closed-set”, that is, a fixed number of predetermined classes selected due to their relevance to the respective study’s subject matter (e.g., Figure 5). It should be noted that such classifiers are only capable of assigning labels from this predetermined set, even in cases where a test sample is not represented in the original set. As a result, when deployed in scenarios where collected data include plankton classes in the original dataset along with non-plankton objects and organisms not included in the original dataset, classifier accuracy and precision underperform test-projected estimates in practice (Eerola et al., 2024; Pu et al., 2021).

To illustrate this point, a collection of particles was annotated but not included in the classifier training dataset. Introducing these OOD objects into the test dataset led to a decrease in accuracy from 94% to 63% (Figure 6A). Curating an exhaustive dataset to represent open environmental systems (e.g., freshwater lakes) is currently impractical due to the labor (sampling and annotation) required. To begin to address the OOD-associated loss in model performance, a bespoke class representing particles outside of the standard set of phytoplankton taxa of interest, referred to as the “known OOD class” (KOC) was introduced to the dataset prior to classifier training with the aim of reducing the classifier’s epistemic uncertainty. In theory, this would refine the decision boundaries associated with original primary target classes, ultimately enhancing overall model performance. To investigate the effect of the KOC class on OOD object recognition, the OOD particles were divided into two categories: debris class, which consists of non-planktonic particles, and novel class, containing plankton species outside of the “known” classes (**Figure S8**). The KOC was strictly limited to the debris class, ensuring that the novel class remained entirely unknown to the model. With the inclusion of the KOC, classifier accuracy increased from 63% to 77% (Figure 6A). Figure 6B-E illustrates a two-dimensional view of the N-dimensional feature space encoded by the classification model using t-SNE (Van Der Maaten and Hinton, 2008), for classifier variants trained without (Figure 6B **and 6C**) and with (Figure 6D **and 6E**) the KOC. Regions are colored as the mapping of features associated with specific classes in the training set, and black scatter points correspond to individual unknown OOD particles. The KOC-trained variant exhibits greater inter-region distances and a higher proportion of unknown OOD particles outside of class-associated regions, i.e., in the “unassigned” space. This behavior is advantageous in that fewer unknown OOD particles were misclassified as a primary target class, and in post-processing these objects can be assigned a null class label independent from the original class labels. In aggregating particles with this label assignment, processed samples can then be reviewed by a human annotator to examine these specific particles.

**Figure 6:**
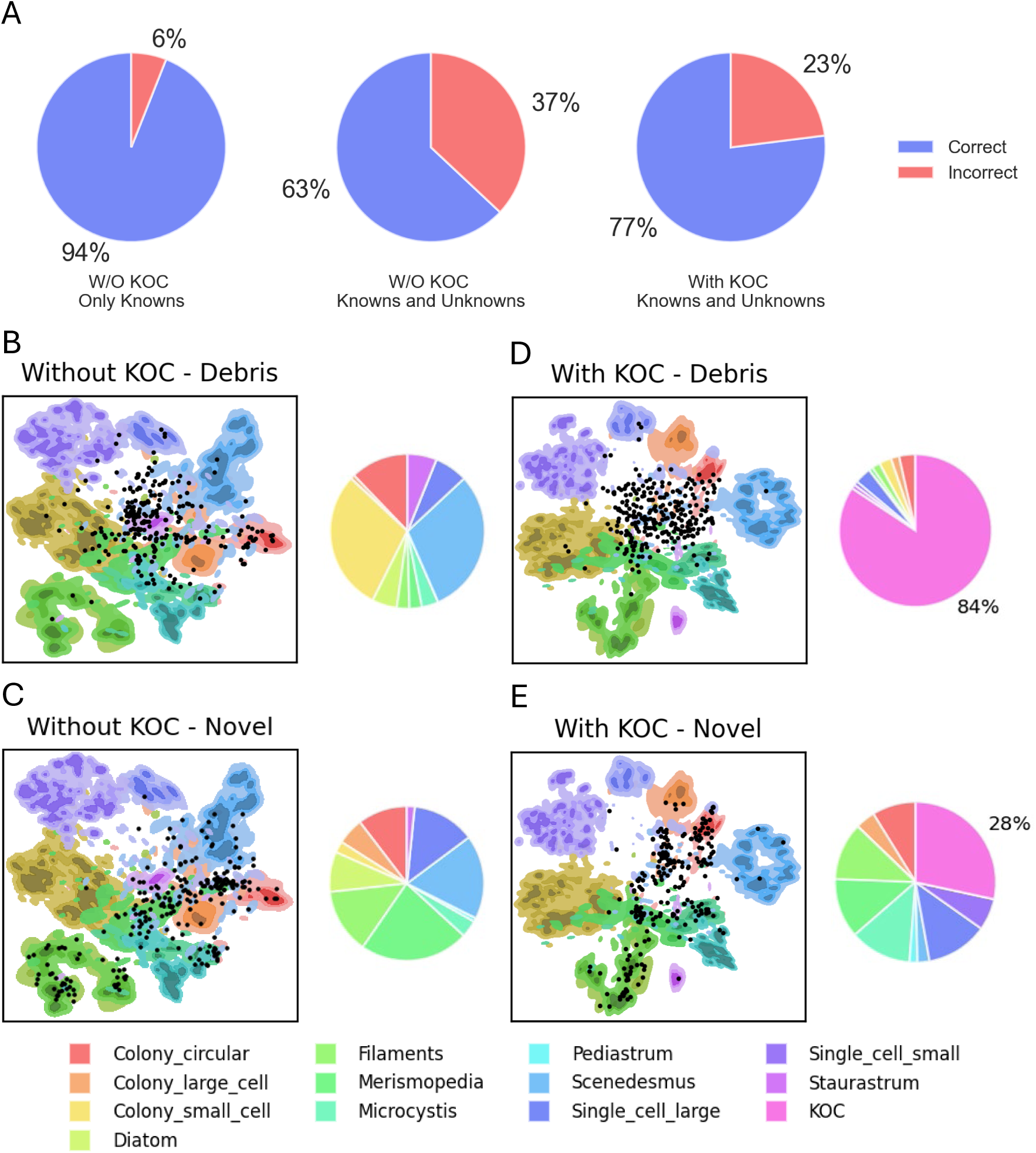
(A) Pie chart illustrating the prediction performance of CNN models trained with and without the KOC class on known and unknown particles. The percentages represent the fractions of correct and incorrect predictions for each model. (B) - (E) Density contour plot for models trained without (B, C) and with (D, E) the KOC, generated from t-SNE projections, with color-coded contour levels for each class in the training dataset. Black points represent the ’debris’ class (B, D) and ‘novel’ class (C, E) from the test set, showing its distribution across different class densities. The accompanying pie chart shows the fraction of the ’debris’ class and ‘novel’ class classified as various classes for each model. The percentages represent the fractions of Debris or Novel class predicted as KOC.

### Classification with rejection using probability thresholding may allow for optimized annotation of novel classes

Even with the inclusion of KOC, eliminating misclassifications is challenging due to the high diversity of planktonic microorganisms and non-cellular debris in samples analyzed from open environmental systems. Given this complexity, phytoplankton monitoring in environmental samples aligns more closely with a “classification with rejection” or “open-set recognition” task where samples contain numerous types of both biotic and abiotic particles, often distinct from the classes the model was trained on. Previous studies (Faillettaz et al., 2016; Luo et al., 2018; Xu et al., 2022) have employed probability thresholding to mitigate misclassification. Probability thresholding involves only retaining predictions with a probability value higher than a given threshold. As the threshold increases, precision tends to increase as label assignment becomes stricter. Figure 7A shows the precision-recall curves for varying probability thresholds, comparing CNN models utilizing SoftMax thresholding both with and without the inclusion of the KOC during the training phase. The addition of KOC to the training set enhances prediction precision when the test set contains both known and unknown classes. Thus, the use of SoftMax thresholding can improve precision but at the cost of recall. For instance, setting a threshold cutoff corresponding to 70% recall for the model trained with KOC (represented by the green curve) results in an overall precision exceeding 82%. Moreover, adjusting the threshold cutoff to achieve 60% recall can elevate precision beyond 90%. This approach may offer operational benefits by consolidating erroneous predictions into a single ‘OOD’ class rather than distributing them across all classes, thereby simplifying manual verification. This increase in precision supports the validity of probability thresholding as an effective method for minimizing erroneous predictions in the autonomous monitoring of algae in freshwater environments.

**Figure 7:**
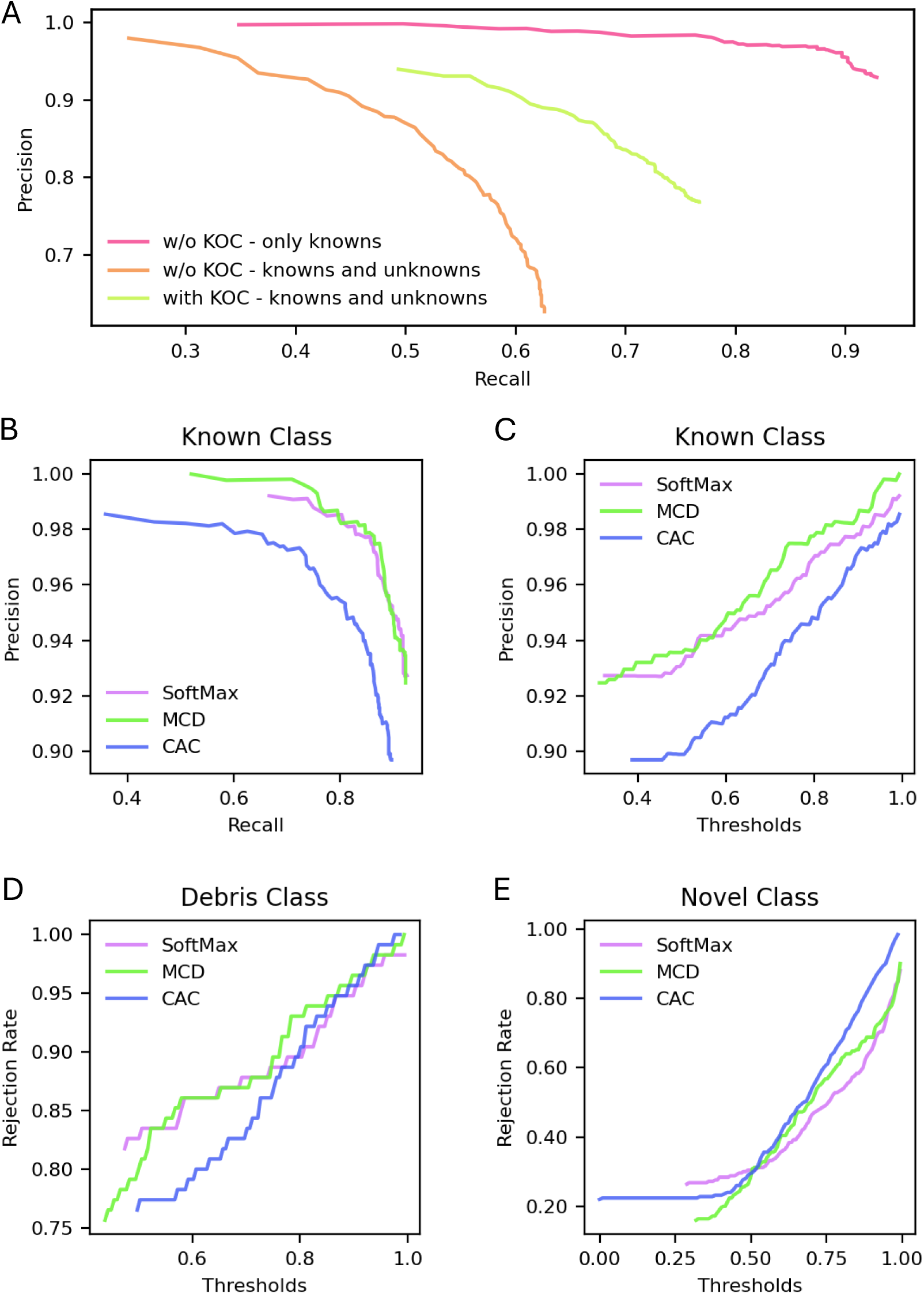
(A) Precision-recall curve for SoftMax models trained with (green) and without (orange, violet) KOC on only known particles (orange) and known and unknown (debris, novel) particles (green, violet). (B) Precision of SoftMax, MCD, and CAC models for varying thresholds in known class identification. (C) Precision-recall curve for the three models on the known class. (D, E) Rejection rate of the three models at varying thresholds for the debris class and novel class, respectively.

However, as the threshold increases, a growing number of predictions are rejected or labeled as OOD, potentially leading to lower detection rates for low-density or rare particles. To further investigate precision and recall in the classification of non-curated datasets, two additional methods for detecting unknown OOD particles were explored: Monte Carlo Dropout (MCD) and Class-Anchored Clustering (CAC). The MCD method involves dropping out certain nodes during inference to generate multiple predictions for the same input, thereby estimating uncertainty by analyzing the variability in these predictions. In contrast, CAC is a distance-based open-set classifier that uses a distance metric to predict the class of a particle. This method is optimized to increase inter-class distance while decreasing intra-class distance (Miller et al., 2021). To understand how rejection through thresholding affects the prediction precision, the analysis was done for known class, debris class, and novel class separately.

Figure 7B illustrates the precision-recall curves for the three models focusing on the known class. Notably, SoftMax and MCD demonstrate a similar relationship between precision and recall, while CAC exhibits lower precision across all recall values. When targeting a 90% recall for the known class, both SoftMax and MCD achieved precision rates of 95%, whereas CAC achieved a lower precision of 90%. This finding underscores the superior performance of SoftMax and MCD in effectively classifying known classes compared to CAC. The thresholds corresponding to this 90% recall rate for the known class are 0.67, 0.6, and 0.45 for SoftMax, MCD, and CAC, respectively (Figure 7C). These thresholds resulted in the rejection of 87%, 86%, and 77% of the debris class (Figure 7D), and 42%, 40%, and 22% of the novel class (Figure 7E) for SoftMax, MCD, and CAC, respectively. For high known class recall (>=90%), SoftMax and MCD can reject a higher percentage of particles in the novel class. Lowering the recall target to 80% further elevates the precision rates for SoftMax and MCD to greater than 98%, with CAC also achieving over 95% precision (Figure 7B). The corresponding thresholds for 80% recall for the known class are 0.94, 0.89, and 0.83 for SoftMax, MCD, and CAC (Figure 7C), leading to the rejection of 97%, 95%, and 93% of the debris class (Figure 7D), as well as only 0.72%, 0.69%, and 0.74% of the novel class for SoftMax, MCD, and CAC, respectively (Figure 7E). For lower known class recall (≤80%), CAC can reject more novel classes than SoftMax and MCD.

In general, CAC demonstrated a higher OOD detection rate and superior performance at lower recall values for the known class, while both CAC and MCD outperformed at higher known class recall levels. This suggests that no single method is definitively superior in all scenarios, and the choice depends on the specific use case. For applications requiring a high OOD identification or rejection rate, CAC would be the preferred method. Conversely, when a high yield of known class predictions is necessary, SoftMax or MCD would be more suitable. However, all three classification-with-rejection methods—CAC, MCD, and SoftMax thresholding—outperformed the closed-set classification approach, highlighting their overall effectiveness in handling OOD particles in environmental monitoring tasks.

### Towards open world recognition for automated classification of phytoplankton in open environmental systems

Our findings suggest that integrating a known OOD class (KOC) into the classifier significantly enhances its ability to detect unknown particles and improves overall model performance. Classifiers typically struggle with OOD predictions due to epistemic uncertainty, often stemming from insufficient training data. By incorporating a known OOD class, this uncertainty is reduced, leading to more refined decision boundaries for known classes. Transitioning from a closed-set classifier to a classification model with a rejection option improves accuracy and precision while reducing misclassification rates. This approach also enables the classification model to identify and isolate particles with high prediction uncertainty, providing an opportunity for manual annotation and subsequent incorporation into the training dataset, thereby increasing the model’s robustness. In practice, future efforts should concentrate on collecting more images of microalgae, phytoplankton, and other particles throughout the deployment lifespan of in-situ monitoring devices like ARTiMiS and utilize classification-with-rejection methods to effectively differentiate between known and OOD particles. These OOD particles can then be categorized using unsupervised clustering methods or semi-supervised classification methods followed by manual curation. Periodically updating the training dataset with newly acquired images and retraining the classifier on this updated dataset will reduce epistemic uncertainty and enhance the classification robustness for environmental monitoring; this approach, known as “Open World Recognition (OWR),” (Figure 8) was first defined by Bendale and Boult, (2015). An OWR system requires three tasks: classifying known and OODs, clustering OODs and labeling them as novel classes, and applying incremental learning (Gao et al., 2024; Joseph et al., 2021). This study suggests that OWR could be the optimal approach for real-time monitoring of freshwater phytoplankton, enabled by low-cost FIM platforms such as ARTiMiS, thus offering a dynamic and robust solution for managing the inherent variability of open environmental systems.

**Figure 8:**
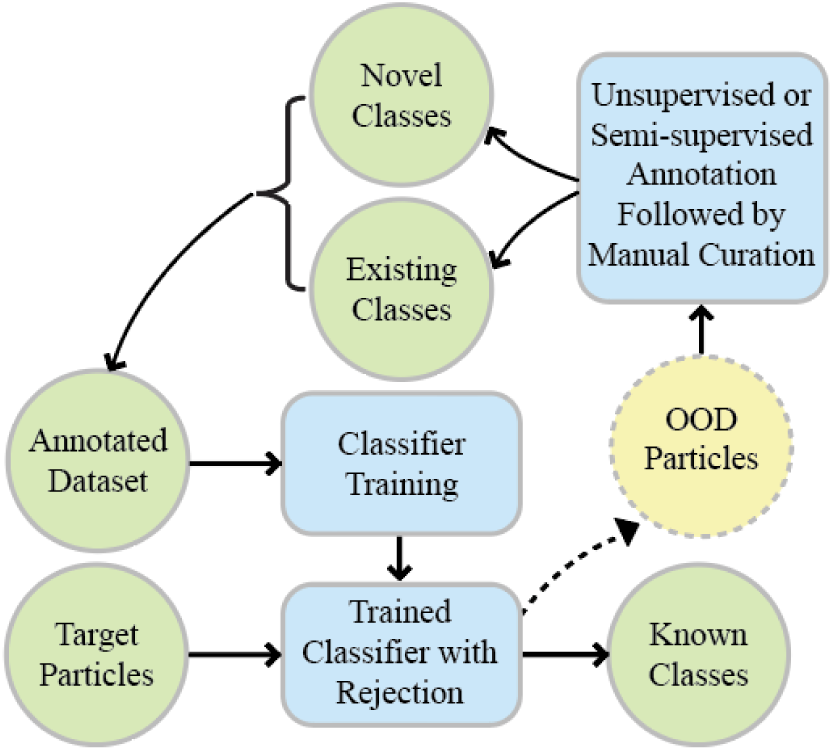
Suggested OWR framework for automated environmental monitoring system. OOD particle images, identified through classification with rejection, require re-annotation and integration into the annotated dataset for continuous improvement of the system

## Conclusions

1. Effective real-time monitoring of microalgae requires robust automated processes to address challenges such as OOF particles, background particles, and OOD particles.
2. A Random Forest model utilizing semantic features demonstrates greater generalizability compared to CNN trained exclusively on image data for OOF particle detection.
3. A two-stage clustering algorithm based on DBSCAN is effective at identifying and removing background particles in flow imaging microscopy.
4. While a closed-set CNN classifier achieves high accuracy for known classes, its performance deteriorates significantly when encountering OOD particles. Therefore, closed-set classification is not ideal for autonomous monitoring of environmental samples.
5. Incorporating a subset of OOD particles into the training dataset refines decision boundaries for classifiers, enhancing the accurate detection of OOD particles.
6. Employing a classification-with-rejection model improves both precision and accuracy when characterizing environmental samples.
7. Implementing an OWR framework, supported by classification with rejection, is crucial for advancing automated environmental monitoring systems.

## Acknowledgments

This research was supported by the Water Research Foundation (Project 5154), the Paul L Busch Award for Applied Water Quality Research, and the National Science Foundation (NSF) (CBET Award Number: 2220792),

## References

Álvarez, E., López-Urrutia, Á., Nogueira, E., 2012. Improvement of plankton biovolume estimates derived from image-based automatic sampling devices: application to FlowCAM. J Plankton Res 34, 454–469. 10.1093/plankt/fbs017

Bendale, A., Boult, T., 2015. Towards Open World Recognition, in: Proceedings of the IEEE Conference on Computer Vision and Pattern Recognition (CVPR). pp. 1893–1902.

Caballero, I., Fernández, R., Escalante, O.M., Mamán, L., Navarro, G., 2020. New capabilities of Sentinel-2A/B satellites combined with in situ data for monitoring small harmful algal blooms in complex coastal waters. Sci Rep 10, 8743. 10.1038/s41598-020-65600-1

Camoying, M.G., Yñiguez, A.T., 2016. FlowCAM optimization: Attaining good quality images for higher taxonomic classification resolution of natural phytoplankton samples. Limnol Oceanogr Methods 14, 305–314. 10.1002/lom3.10090

Cen, C., Zhang, K., Zhang, T., Zhou, X., Pan, R., 2020. Algae-induced taste and odour problems at low temperatures and the cold stress response hypothesis. Appl Microbiol Biotechnol 104, 9079–9093. 10.1007/s00253-020-10884-6

Chan, W.H., Fung, B.S.B., Tsang, D.H.K., Lo, I.M.C., 2023. A freshwater algae classification system based on machine learning with StyleGAN2-ADA augmentation for limited and imbalanced datasets. Water Res 243, 120409. 10.1016/j.watres.2023.120409

Choran, N., Örmeci, B., 2023. Micro-flow imaging for in-situ and real-time enumeration and identification of microplastics in water. Frontiers in Water 5, 1148379. 10.3389/frwa.2023.1148379

Ciranni, M., Murino, V., Odone, F., Pastore, V.P., 2024. Computer vision and deep learning meet plankton: Milestones and future directions. Image Vis Comput 143, 104934. 10.1016/j.imavis.2024.104934

Correa, I., Drews, P., Botelho, S., De Souza, M.S., Tavano, V.M., 2017. Deep learning for microalgae classification, in: Proceedings - 16th IEEE International Conference on Machine Learning and Applications, ICMLA 2017. Institute of Electrical and Electronics Engineers Inc., pp. 20–25. 10.1109/ICMLA.2017.0-183

Deglint, J.L., Tang, L., Wang, Y., Jin, C., Wong, A., 2018. SAMSON: Spectral Absorption- fluorescence Microscopy System for ON-site-imaging of algae. Journal of Computational Vision and Imaging Systems 4(1), 3. 10.15353/jcvis.v4i1.324

Den Uyl, P.A., Thompson, L.R., Errera, R.M., Birch, J.M., Preston, C.M., Ussler, W., Yancey, C.E., Chaganti, S.R., Ruberg, S.A., Doucette, G.J., Dick, G.J., Scholin, C.A., Goodwin, K.D., 2022. Lake Erie field trials to advance autonomous monitoring of cyanobacterial harmful algal blooms. Front Mar Sci 9, 1021952. 10.3389/fmars.2022.1021952

Dodds, W.K., Bouska, W.W., Eitzmann, J.L., Pilger, T.J., Pitts, K.L., Riley, A.J., Schloesser, J.T., Thornbrugh, D.J., 2009. Eutrophication of U.S. Freshwaters: Analysis of Potential Economic Damages. Environ Sci Technol 43, 12–19. 10.1021/es801217q

Eerola, T., Batrakhanov, D., Barazandeh, N.V., Kraft, K., Haraguchi, L., Lensu, L., Suikkanen, S., Seppälä, J., Tamminen, T., Kälviäinen, H., 2024. Survey of automatic plankton image recognition: challenges, existing solutions and future perspectives. Artif Intell Rev 57, 114. 10.1007/s10462-024-10745-y

Ester, M., Kriegel, H.-P., Sander, J., Xu, X., 1996. A density-based algorithm for discovering clusters in large spatial databases with noise, in: Proceedings of the Second International Conference on Knowledge Discovery and Data Mining, KDD’96. AAAI Press, pp. 226– 231.

Faillettaz, R., Picheral, M., Luo, J.Y., Guigand, C., Cowen, R.K., Irisson, J.-O., 2016. Imperfect automatic image classification successfully describes plankton distribution patterns. Methods in Oceanography 15–16, 60–77. 10.1016/j.mio.2016.04.003

Fragoso, G.M., Poulton, A.J., Pratt, N.J., Johnsen, G., Purdie, D.A., 2019. Trait-based analysis of subpolar North Atlantic phytoplankton and plastidic ciliate communities using automated flow cytometer. Limnol Oceanogr 64, 1763–1778. 10.1002/lno.11189

Gao, F., Zhong, W., Cao, Z., Peng, X., Li, Z., 2024. Opengcd: Assisting Open World Recognition with Generalized Category Discovery. 10.2139/ssrn.4903867

Gincley, B., Khan, F., Hartnett, E., Fisher, A., Pinto, A.J., 2024. Introducing ARTiMiS: A Low-Cost Flow Imaging Microscope for Microalgal Monitoring. Environ Sci Technol 58, 13540– 13551. 10.1021/acs.est.4c01928

Hardison, D.R., Holland, W.C., Currier, R.D., Kirkpatrick, B., Stumpf, R., Fanara, T., Burris, D., Reich, A., Kirkpatrick, G.J., Litaker, R.W., 2019. HABscope: A tool for use by citizen scientists to facilitate early warning of respiratory irritation caused by toxic blooms of Karenia brevis. PLoS One 14, e0218489. 10.1371/journal.pone.0218489

Heidi M. Sosik, Emily E. Peacock, Emily F. Brownlee, 2015. WHOI-Plankton, Annotated Plankton Images - Data Set for Developing and Evaluating Classification Methods. URL http://hdl.handle.net/10.1575/1912/7341. (accessed 8.24.24).

Heil, C.A., Muni-Morgan, A.L., 2021. Florida’s Harmful Algal Bloom (HAB) Problem: Escalating Risks to Human, Environmental and Economic Health With Climate Change. Front Ecol Evol 9, 299. 10.3389/fevo.2021.646080

Jetoo, S., Grover, V.I., Krantzberg, G., 2015. The toledo drinking water advisory: Suggested application of the water safety planning approach. Sustainability (Switzerland) 7, 9787– 9808. 10.3390/su7089787

Joseph, K.J., Khan, S., Khan, F.S., Balasubramanian, V.N., 2021. Towards Open World Object Detection, in: Proceedings of the IEEE/CVF Conference on Computer Vision and Pattern Recognition (CVPR). pp. 5830–5840.

Kim, H., Gerber, L.C., Chiu, D., Lee, S.A., Cira, N.J., Xia, S.Y., Riedel-Kruse, I.H., 2016. LudusScope: Accessible Interactive Smartphone Microscopy for Life-Science Education. PLoS One 11, e0162602. 10.1371/journal.pone.0162602

Kramer, B.J., Davis, T.W., Meyer, K.A., Rosen, B.H., Goleski, J.A., Dick, G.J., Oh, G., Gobler, C.J., 2018. Nitrogen limitation, toxin synthesis potential, and toxicity of cyanobacterial populations in Lake Okeechobee and the St. Lucie River Estuary, Florida, during the 2016 state of emergency event. PLoS One 13. 10.1371/journal.pone.0196278

Lumini, A., Nanni, L., 2019. Deep learning and transfer learning features for plankton classification. Ecol Inform 51, 33–43. 10.1016/j.ecoinf.2019.02.007

Luo, J.Y., Irisson, J., Graham, B., Guigand, C., Sarafraz, A., Mader, C., Cowen, R.K., 2018. Automated plankton image analysis using convolutional neural networks. Limnol Oceanogr Methods 16, 814–827. 10.1002/lom3.10285

Marvin, L., Paiva, W., Gill, N., Morales, M.A., Halpern, J.M., Vesenka, J., Balog, E.R.M., 2019. Flow imaging microscopy as a novel tool for high-throughput evaluation of elastin-like polymer coacervates. PLoS One 14, e0216406. 10.1371/journal.pone.0216406

Miller, D., Sunderhauf, N., Milford, M., Dayoub, F., 2021. Class Anchor Clustering: A Loss for Distance-based Open Set Recognition, in: 2021 IEEE Winter Conference on Applications of Computer Vision (WACV). IEEE, pp. 3569–3577. 10.1109/WACV48630.2021.00361

Olsen, M.G., Adrian, R.J., 2000. Out-of-focus effects on particle image visibility and correlation in microscopic particle image velocimetry. Exp Fluids 29, S166–S174. 10.1007/s003480070018

Owen, B.M., Hallett, C.S., Cosgrove, J.J., Tweedley, J.R., Moheimani, N.R., 2022. Reporting of methods for automated devices: A systematic review and recommendation for studies using FlowCam for phytoplankton. Limnol Oceanogr Methods 20, 400–427. 10.1002/LOM3.10496

Pollina, T., Larson, A.G., Lombard, F., Li, H., Le Guen, D., Colin, S., de Vargas, C., Prakash, M., 2022. PlanktoScope: Affordable Modular Quantitative Imaging Platform for Citizen Oceanography. Front Mar Sci 9, 949428. 10.3389/fmars.2022.949428

Pu, Y., Feng, Z., Wang, Z., Yang, Z., Li, J., 2021. Anomaly Detection for in situ Marine Plankton Images. Proceedings of the IEEE International Conference on Computer Vision 2021- October, 3654–3664. 10.1109/ICCVW54120.2021.00409

Richardson, T.L., Lawrenz, E., Pinckney, J.L., Guajardo, R.C., Walker, E.A., Paerl, H.W., MacIntyre, H.L., 2010. Spectral fluorometric characterization of phytoplankton community composition using the Algae Online Analyser®. Water Res 44, 2461–2472. 10.1016/j.watres.2010.01.012

Romero-Martínez, L., van Slooten, C., Nebot, E., Acevedo-Merino, A., Peperzak, L., 2017. Assessment of imaging-in-flow system (FlowCAM) for systematic ballast water management. Science of The Total Environment 603–604, 550–561. 10.1016/j.scitotenv.2017.06.070

Sanseverino, I., António, D.C., Loos, R., Lettieri, T., 2017. Cyanotoxins: methods and approaches for their analysis and detection. Joint Research Centre (JRC), European Union. 10.2760/36186

Sediq, A.S., Klem, R., Nejadnik, M.R., Meij, P., Jiskoot, W., 2018. Label-Free, Flow-Imaging Methods for Determination of Cell Concentration and Viability. Pharm Res 35, 150. 10.1007/s11095-018-2422-5

Sieracki, C.K., Sieracki, M.E., Yentsch, C.S., 1998. An imaging-in-flow system for automated analysis of marine microplankton. Mar Ecol Prog Ser 168, 285–296. 10.3354/MEPS168285

Smith, R.B., Bass, B., Sawyer, D., Depew, D., Watson, S.B., 2019. Estimating the economic costs of algal blooms in the Canadian Lake Erie Basin. Harmful Algae 87, 101624. 10.1016/j.hal.2019.101624

Srivastava, N., Hinton, G., Krizhevsky, A., Salakhutdinov, R., 2014. Dropout: A Simple Way to Prevent Neural Networks from Overfitting. Journal of Machine Learning Research 15, 1929–1958.

Stauffer, B.A., Bowers, H.A., Buckley, E., Davis, T.W., Johengen, T.H., Kudela, R., McManus, M.A., Purcell, H., Smith, G.J., Vander Woude, A., Tamburri, M.N., 2019. Considerations in Harmful Algal Bloom Research and Monitoring: Perspectives From a Consensus-Building Workshop and Technology Testing. Front Mar Sci 6, 460315. 10.3389/fmars.2019.00399

Stroming, S., Robertson, M., Mabee, B., Kuwayama, Y., Schaeffer, B., 2020. Quantifying the Human Health Benefits of Using Satellite Information to Detect Cyanobacterial Harmful Algal Blooms and Manage Recreational Advisories in U.S. Lakes. Geohealth 4, e2020GH000254. 10.1029/2020GH000254

Toldrà, A., O’Sullivan, C.K., Campàs, M., 2019. Detecting Harmful Algal Blooms with Isothermal Molecular Strategies. Trends Biotechnol 37, 1278–1281. 10.1016/j.tibtech.2019.07.003

Van Der Maaten, L., Hinton, G., 2008. Visualizing Data using t-SNE. Journal of Machine Learning Research 9, 2579–2605.

Watson, S.B., Ridal, J., Boyer, G.L., 2008. Taste and odour and cyanobacterial toxins: Impairment, prediction, and management in the Great Lakes. Canadian Journal of Fisheries and Aquatic Sciences 65, 1779–1796. 10.1139/F08-084

Xu, Linquan, Xu, Linji, Chen, Y., Zhang, Y., Yang, J., 2022. Accurate Classification of Algae Using Deep Convolutional Neural Network with a Small Database. ACS ES&T Water 2, 1921–1928. 10.1021/acsestwater.1c00466

Zhang, Z., Qin, W., Cheng, S., Xu, L., Wang, T., Zhang, X.-X., Wu, B., Yang, L., 2011. Assessing the toxicity of ingested Taihu Lake water on mice via hepatic histopathology and matrix metalloproteinase expression. Ecotoxicology 20, 1047–1056. 10.1007/s10646-011-0617-1

Zölls, S., Weinbuch, D., Wiggenhorn, M., Winter, G., Friess, W., Jiskoot, W., Hawe, A., 2013. Flow Imaging Microscopy for Protein Particle Analysis—A Comparative Evaluation of Four Different Analytical Instruments. AAPS J 15, 1200. 10.1208/S12248-013-9522-2

