## Supplemental Information for "Integrating Machine Learning with Flow-Imaging Microscopy for Automated Monitoring of Algal Blooms"

**Table S1:** Features measured during semantic feature extraction. Features that are derived entirely or predominantly from an external computer vision library are indicated by dependency source. Features without a cited library dependency are derived using mathematical equations or methods from the Python standard library.

| Feature Name | Description | Library Dependency | Feature used for RF | Feature used for BPD |
|---|---|---|---|---|
| UUID | Universally Unique Identifier, unique for each particle. | standard | No | No |
| XY Coordinates | Image coordinates of center of crop region. | standard | No | Yes |
| Area | Sum of count of pixels comprising dominant object in foreground. | scikit-image | No | Yes |
| Bounding Box Area | Area of the smallest rectangle enclosing the dominant object. | scikit-image | No | Yes |
| Convex Area | Area of the convex hull of the dominant object. | scikit-image | No | Yes |
| Total Area | Sum of all pixels in binarSy image foreground (can include other objects). | standard | No | Yes |
| Patch Size | Size of the image crop (ROI), in pixels and microns. | standard | No | Yes |
| Mean Intensity | Mean pixel brightness value across entire ROI. | scikit-image | Yes | Yes |
| Maximum Intensity | Maximum pixel brightness value across entire ROI. | standard | Yes | Yes |
| Object Intensity Sum | Sum of pixel brightness values for all pixels comprising the dominant object. | standard | No | Yes |
| Total Intensity Sum | Sum of all pixel brightness values across the entire ROI. | standard | No | Yes |
| Solidity | Ratio of pixels in the enclosing region to pixels of the convex hull image. | scikit-image | Yes | Yes |
| Perimeter | Perimeter contour calculated through the centers of border pixels using a 4-connectivity. | scikit-image | No | Yes |
| Equivalent Diameter | Diameter of a circle with the same area as the region. | scikit-image | No | Yes |
| Feret Diameter (max) | Maximum Feret's diameter, the longest distance between points around the convex hull. | scikit-image | No | Yes |
| Major Axis Length | Length of major axis of an ellipse having the same normalized second central moments as the dominant object. | scikit-image | No | Yes |
| Minor Axis Length | Length of minor axis of an ellipse having the same normalized second central moments as the dominant object. | scikit-image | Yes | Yes |
| Eccentricity | Ratio of the focal distance over the major axis length, calculated from an ellipse having the same normalized second central moments as the dominant object. | scikit-image | No | Yes |
| Orientation | Angle between the 0th axis and major axis of the ellipse having the same normalized second central moments as the dominant object. | scikit-image | No | Yes |

| | | | | |
|---|---|---|---|---|
| Centroid | Weighted centroid of the image. | scikit-image | No | Yes |
| Hu Moments | Image moments: translation, scale, and rotation invariant. | scikit-image | Yes | Yes |
| Hu Circularity | Circularity of object using the 0th Hu's moment as the object's radius. | scikit-image | No | Yes |
| Entropy | Shannon's entropy of the image. | scikit-image | No | Yes |
| Circularity | Circularity of object using the object's calculated area and perimeter. | standard | No | Yes |
| Euler Number | Euler characteristic of binary image. | scikit-image | No | Yes |
| Object Topography | Distance transform of object, binned as topographic levels. | scipy | No | Yes |
| Aspect Ratio | Ratio of minor axis length to major axis length. | standard | No | Yes |
| Biovolume (Sphere) | Calculated volume of object, assuming sphere (circular) geometry. | standard | No | Yes |
| Biovolume (Spheroid) | Calculated volume of object, assuming spheroid (elliptical) geometry. | standard | Mo | Yes |
| Edge Noise (Laplacian) | Blurriness of object as calculated as the standard deviation of Laplacian of Gaussian blur. | OpenCV | Yes | Yes |
| Edge Noise (Gradient) | Blurriness of object as calculated as the standard deviation of Sobel gradient of Gaussian blur. | OpenCV | Yes | Yes |
| Edge Gradient | Visual gradient at edge of object, calculated as average pixel intensity of object's outer edge. | standard | Yes | Yes |
| Edge Difference | Visual gradient at edge of object, calculated as difference in pixel values outside and inside the object's edge. | standard | Yes | Yes |

**Table S2:** Class labels and corresponding number of image crops

| Class | Number of Images |
|---|---|
| Colony circular | 216 |
| Colony large cell | 304 |
| Colony small cell | 1066 |
| Debris | 760 |
| Diatom | 236 |
| Filaments | 466 |
| Merismopedia | 222 |
| Microcystis | 237 |
| Pediastrum | 285 |
| Scenedesmus | 774 |
| Single cell large | 306 |
| Single cell small | 831 |
| Staurastrum | 68 |

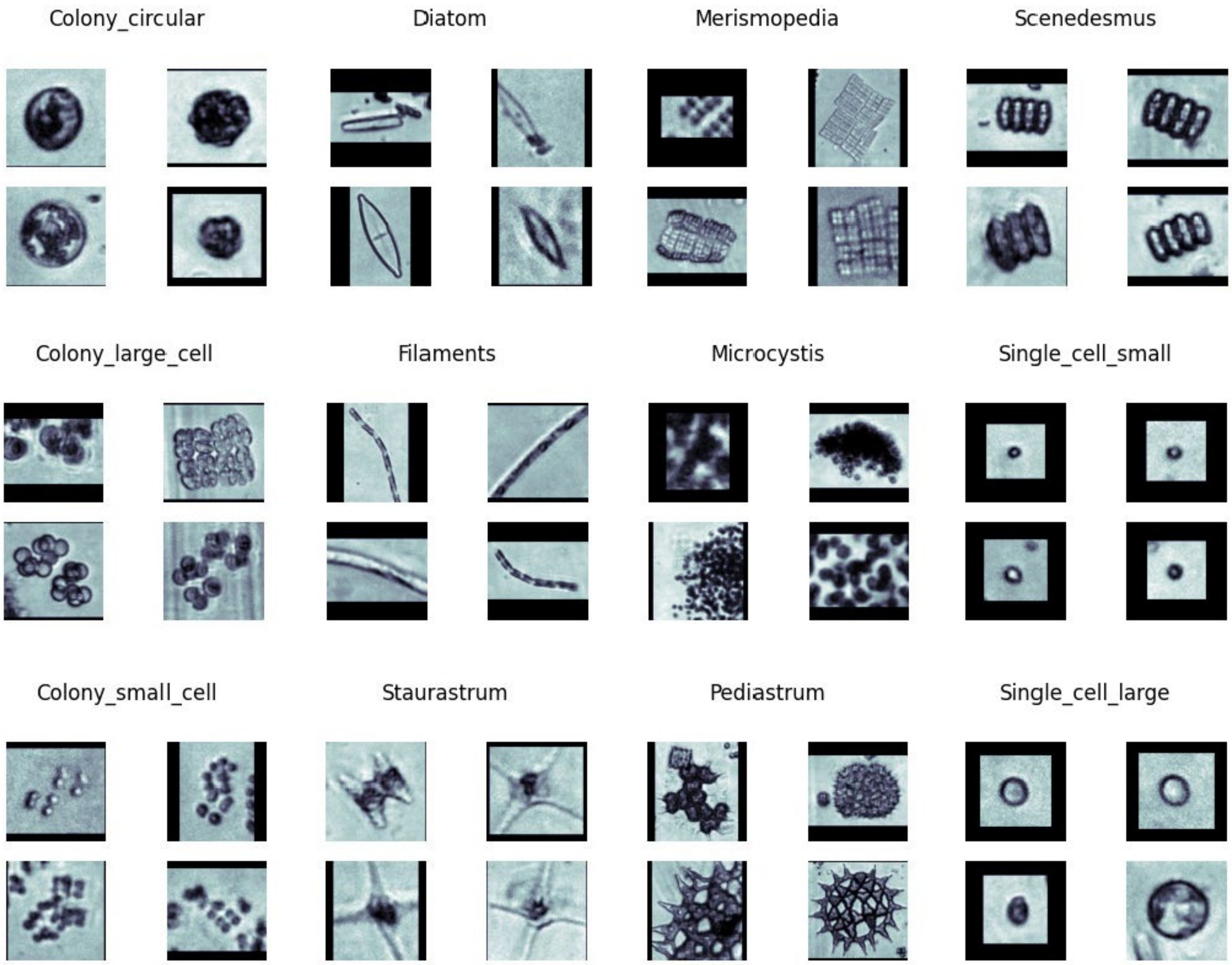


**Figure S1:** Four example images for each of the 12 known classes

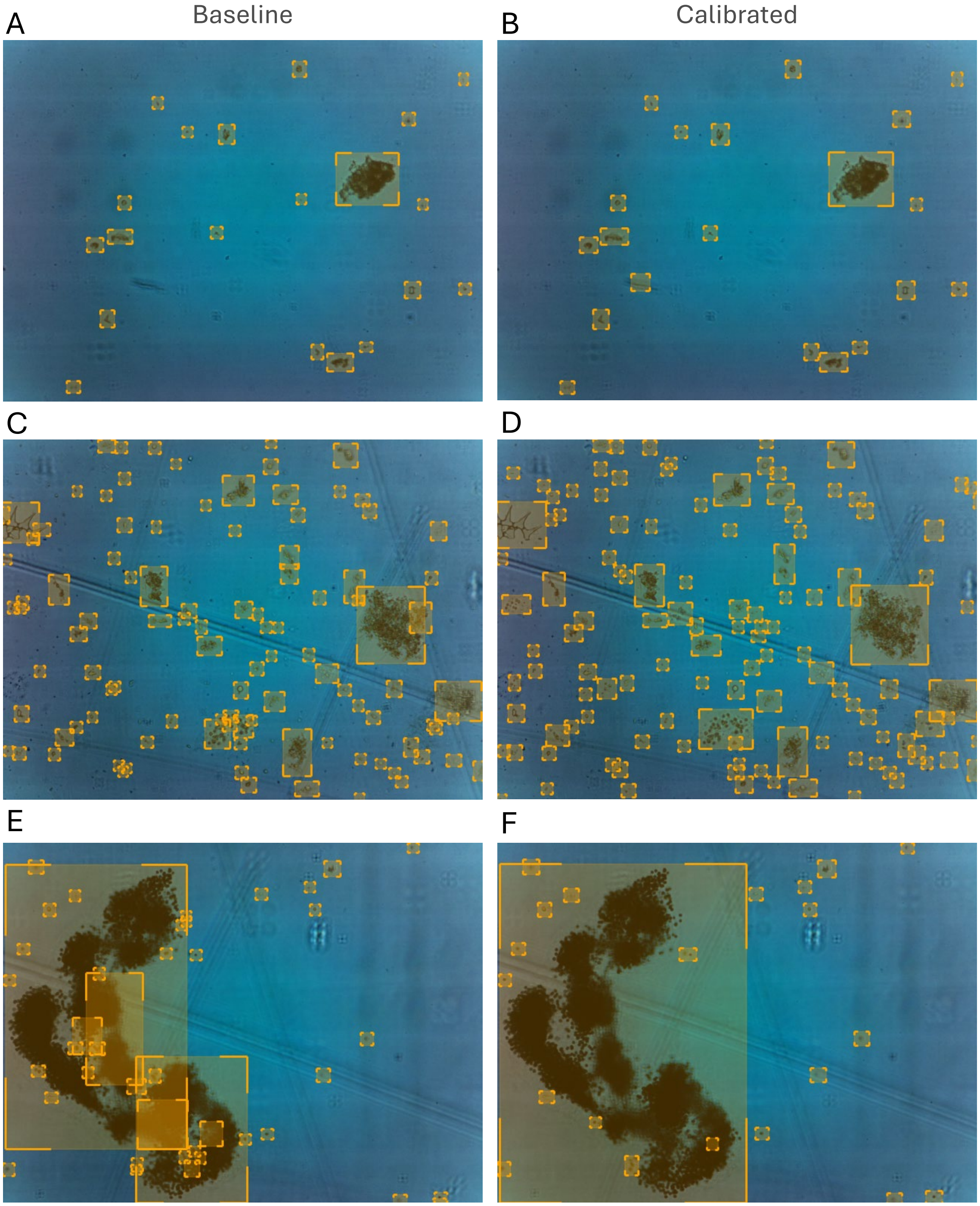


**Figure S2:** Objects detected in widefield images by baseline (A, C, E) and calibrated (B, D, F) ODA. Yellow boxes represent individual objects (either single cells or colonies). (A, B) show image samples for freshwater sparse samples, (C, D) represent dense samples, and (E, F) display image samples for Microcystis colonies.

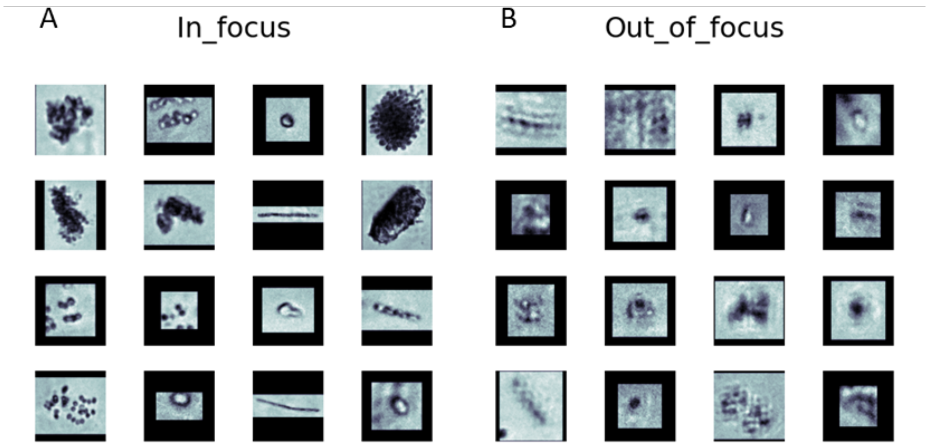


**Figure S3:** Examples of A) in focus and B) out of focus particles

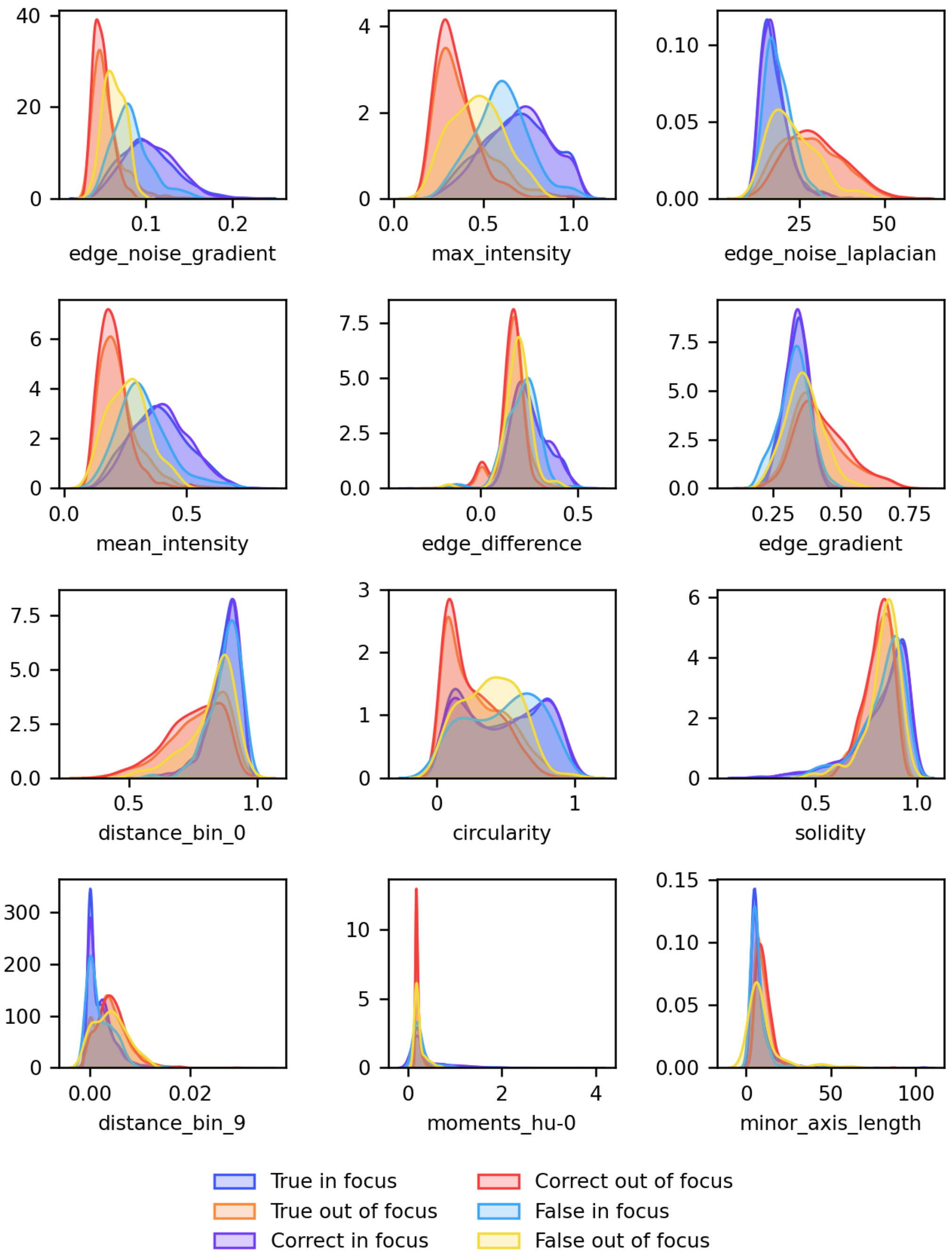


**Figure S4:** Density plot of top 12 features for true class, correct and incorrect predictions

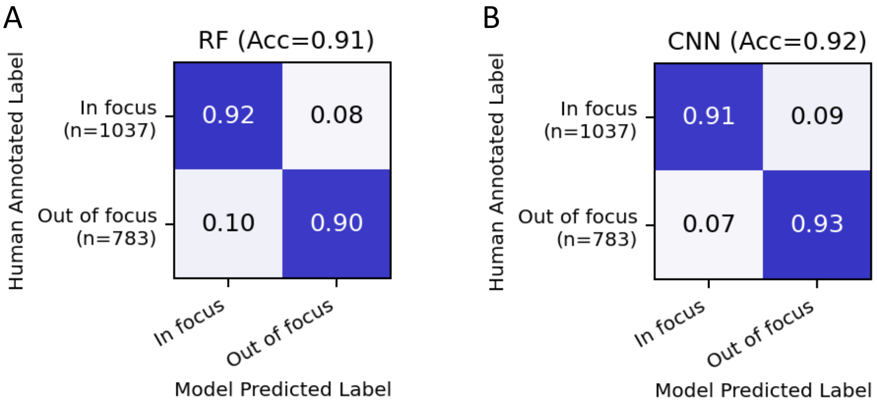


**Figure S5:** Confusion matrices showing performance of A) random forest, B) CNN based OOF detection

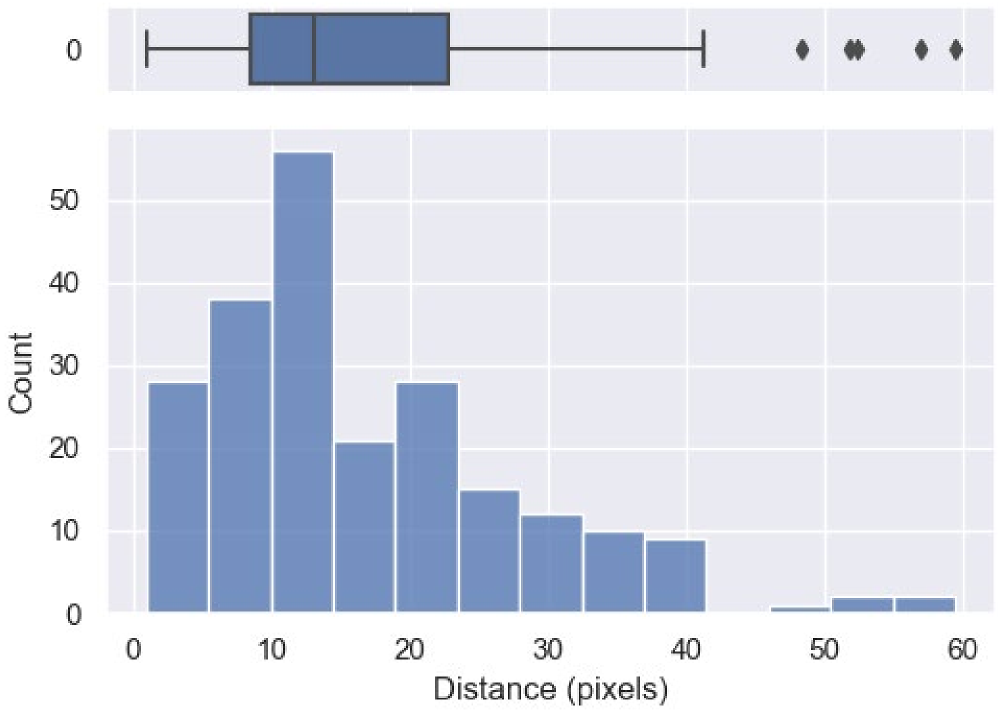


**Figure S6:** Histogram of distances traveled by background particles in consecutive images

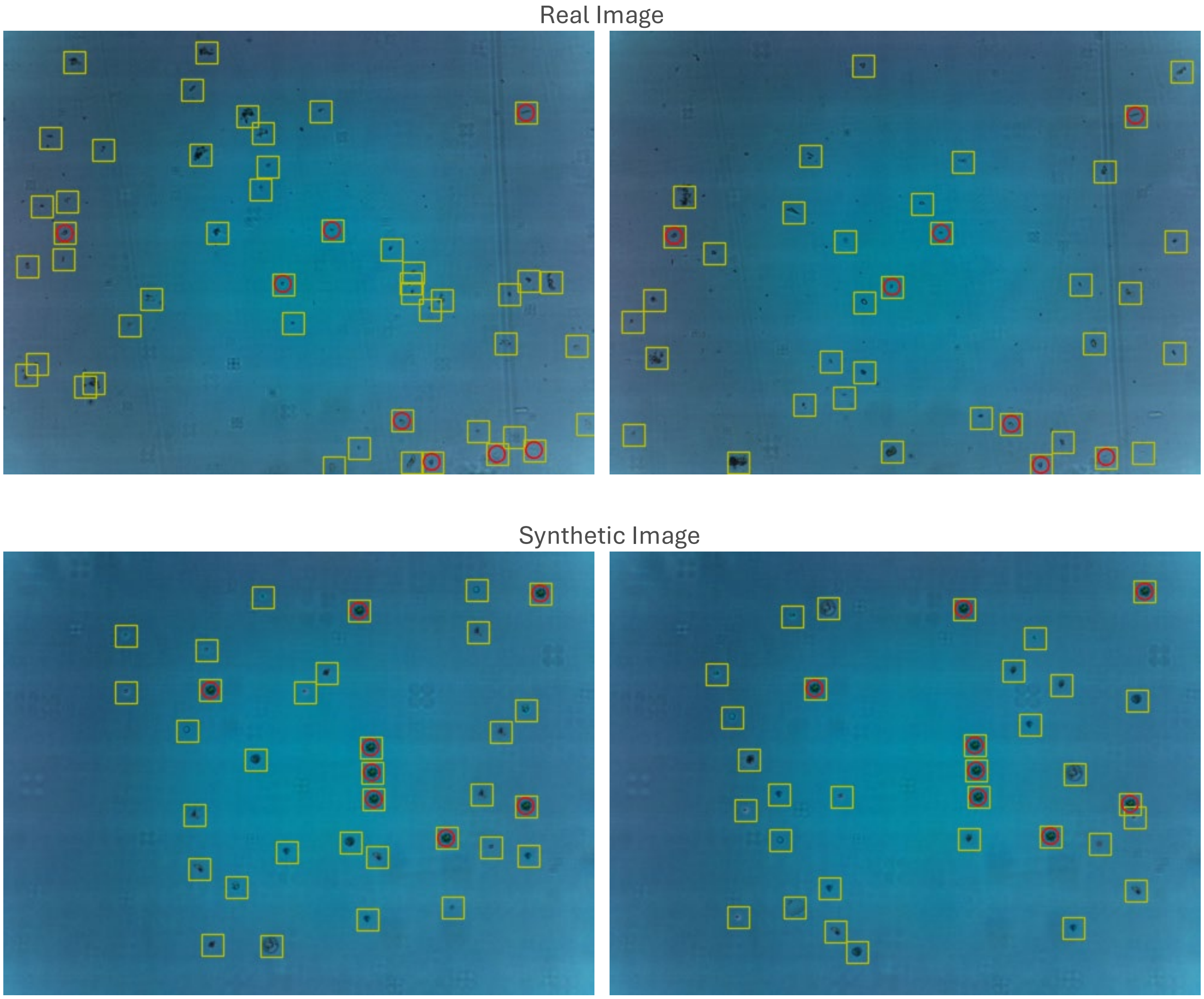


**Figure S7:** Background particle detection by BPD algorithm on real images and synthetic images (yellow box represents particle detected by object detector and red box represents particles identified as background particle in two consecutive images)

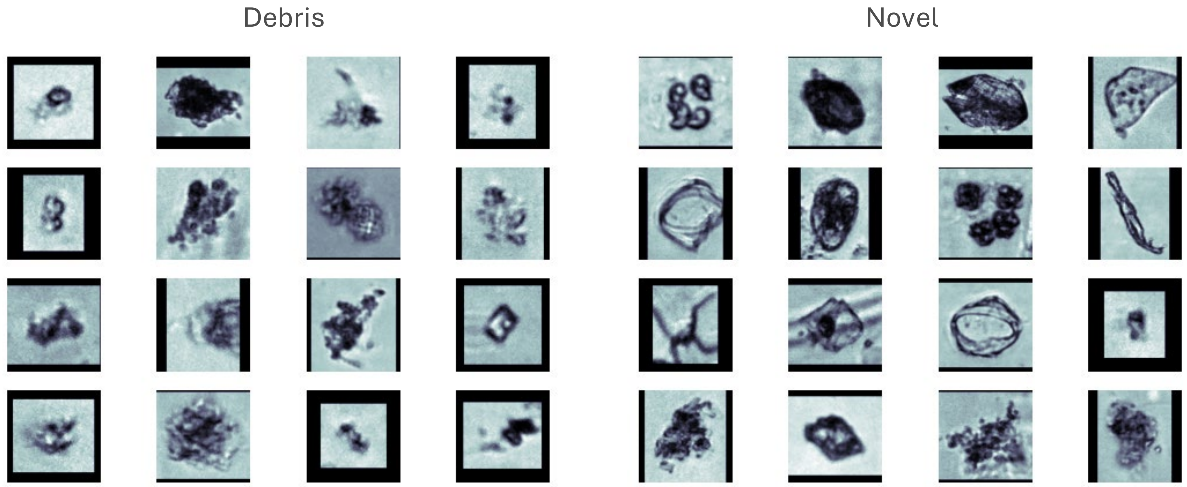


**Figure S8:** Example images OOD particles: debris on the left and novel on the right
